# Developments in the European parasitoid community of *Dryocosmus kuriphilus*

**DOI:** 10.64898/2026.09.01.748543

**Authors:** K. McCormack, J. Cottrell, G. Stone, K. Schönrogge

## Abstract

The invasive gallwasp *Dryocosmus kuriphilus* was first detected in Italy in 2002, although likely to have initially arrived in the late 90s. Its ability to utilise sweet chestnut species non-native to its original Chinese range has allowed it to spread rapidly, and throughout Europe via European sweet chestnut *Castanea sativa*. Given the severity of its impact on *C. sativa* crop production, particularly in Mediterranean countries, previous studies have aimed to assess damage levels caused by *D. kuriphilus,* the efficacy of and potential non-target effects of the introduced biocontrol agent *Torymus sinensis,* and the possibility of regulation by native parasitoids. As yet a broad overview and analytical synthesis of these native parasitoid communities are absent. This review focuses on important aspects of the *D. kuriphilus* invasion. In particular, the invasion history and currently known distribution of *D. kuriphilus*, and several aspects of its associated parasitoid community. For native species to plausibly supress *D. kuriphilus*, we might expect rates of parasitoid attack to increase with establishment time as native populations adapt to exploit the new resource, and this is a key focus of the review. We answer the following questions:

1) What is the distribution of *D. kuriphilus* in Europe, and has *D. kuriphilus* fully utilised the available niche space within its 20+ years in Europe?
2) Which species of native parasitoids attack *D. kuriphilus* in Europe and what are their ecological characteristics?
3) How consistent is the parasitoid community of *D. kuriphilus* across its range, and are there signs of convergence over time?
4) What effect does establishment time have on the species richness and abundance of parasitoid communities?

We report the following:

1) *D. kuriphilus* has expanded its range throughout Europe and is present in nearly every major region where sweet chestnut is present. Native and non-native naturalised chestnut forests may be less susceptible to invasion than areas of industry due to differing socioeconomic and ecological factors, though areas with large chestnut industries also tend to be in the most heavily forested areas in the non-native range of sweet chestnut. *D. kuriphilus* has reportedly been eradicated from some countries, and effectively eradicated in a number of countries implementing biocontrol with *T. sinensis,* although successive invasions from neighbouring regions are still possible, and eradication may be transient.
2) 72 parasitoid species are identified attacking *D. kuriphilus* in Europe (far more species than any other gallwasp in the Western Palearctic). Its members are predominantly oak gallwasp parasitoids (82% of species), followed by gall-specialists of different host plants, leaf miner parasitoids and a minority of others with differing host life stages and ecologies. Parasitoids attacking *D. kuriphilus* are dominated by idiobiont ectoparasitoids of the superfamily Chalcidoidea (>96%).
3) The parasitoid community is highly variable, both temporally and spatially, although the vast proportion (>95%) of parasitoids at any one time are composed of locally common generalist oak gall parasitoids. The most common members include *Bootanomyia dorsalis, Eupelmus urozonus, Eurytoma brunniventris, Mesopolobus sericeus* and *Torymus flavipes*.
4) The length of establishment time has minimal effect on the species richness, abundance, and composition of the community, suggesting that regulation by natives, if it occurs, may take longer than the 20+ years that *D. kuriphilus* has persisted.

While little evidence of increasing parasitoid attack of *D. kuriphilus* is apparent, we exercise caution by stating that the heterogeneity in available data are large, and that common biocontrol interventions using *T. sinensis* interrupt the natural process of community development dramatically. *D. kuriphilus* has been present in Europe for nearly three decades and few localities have repeated years of data collection. Even fewer studies have communities with establishment times exceeding ten years. Proper biocontrol by natives may not occur within short timeframes, although studies of other gallwasp invaders find similar results over periods exceeding 40 years. Given that many countries have chosen to implement *T. sinensis* for biocontrol, the focus may be better spent monitoring native gall communities for potential non-target effects.

## Introduction

*D. kuriphilus* (Cynipinae: Cynipini) is a gallwasp species originally from China, with an invasion history spanning the 1940s to the present day. Initial invasions occurred in Japan and Korea in the 40s (Murakami *et. al.*, 1980; Cho and Lee, 1963), the US in the 70s (Payne *et. al.*, 1976), Nepal in the 90s (Abe *et. al.*, 2007), and Europe via the Calabria region of Northern Italy where it is believed to have arrived in the late 1990s via trade of infected plant material from China (Aebi *et. al.*, 2006; Brussino *et. al..*, 2002). Every country that it has invaded has subsequently been severely impacted by its establishment due to the severity and rapidity of damage to sweet chestnut trees (Kato and Hijii, 1997; Payne *et. al.*, 1983). *D. kuriphilus* is capable of significantly decreasing wood production, and decreasing fruiting output by 40 to 70%, far beyond tolerable levels of industry (Graziosi and Santi, 2008). Several Mediterranean countries in Europe have a long history of chestnut cultivation and chestnut crops are a significant agricultural product, particularly in Italy, France, Spain, Turkey, and Portugal. The European Plant Protection Organization (EPPO) assessed the potential impact of *D. kuriphilus* and the European Food Safety Authority (EFSA) concluded that interventions were necessary (EFSA, 2010).

Several of *D. kuriphilus*’ life history traits likely aid in its ease of spread. Firstly, its reproduction is asexual, circumventing the need for mate-finding during establishment. A single female is capable of ovipositing over 200 eggs (50-100 on average) (EFSA, 2010; Tokuhisa, 1982) and potentially establishing a population. In Europe, sweet chestnut has no native gallwasp community and hence *D. kuriphilus* has no competition for host plant resources. With no native gallwasp community surrounding sweet chestnut as a host, and with a general rule that gallwasps are attacked by gallwasp-specific parasitoids, *D. kuriphilus* theoretically lives within enemy-free space, allowing it to proliferate unabated (although all evidence shows that it is attacked-see below). The most susceptible places to *D. kuriphilus* invasion are chestnut orchards within regions with a high number of degree days, and the generally low genetic diversity of *C. sativa* cultivars and low generational turnaround also likely aid in the rapid spread of and limited resistance to *D. kuriphilus* (EFSA, 2010). Two of the greatest threats to the chestnut industry come from *D. kuriphilus,* and the chestnut blight fungus *Cryphonectria parasitica* (EFSA, 2010). Both species are Asian in origin, and both have been spread to the US, Asia and Europe via trade of infected chestnut material (EFSA 2016; 2010). Within the US, *C. parasitica* has effectively eradicated native US chestnut (*C. dentata*) in several regions, resulting in enormous ecological and economic impacts (Elliot and Swank, 2008). While its impacts are currently less severe in Europe, moderate local impacts are apparent (Rigling and Prospero, 2018). A secondary concern of the spread of *D. kuriphilus* is the possibility that it facilitates transmission of *C. parasitica*, and vice-versa (Pérez-Sierra *et. al.*, 2020; Morales-Rodriguez *et. al.*, 2019; Michaelakis *et. al.*, 2016; Prospero and Forster, 2011). Numerous *Cryphonectria* species including *C. parasitica* have been detected in old galls of *D. kuriphilus* (Pérez-Sierra *et. al.*, 2020; Prospero and Forster, 2011), though not within the adult insects themselves (Morales-Rodriguez *et. al.*, 2019). Hence, while the infection of trees via gall damage is plausible and likely (Meyer *et. al.*, 2015), the gallwasp adults themselves are not implicated propagation vectors.

### Gallwasp invaders as hosts

A common thread amongst most gallwasp invaders throughout Europe has been in their 1) relatively contemporary shared evolutionary history, 2) the general contiguity of their native and invaded ranges, and 3) the common utilisation of oak (*Quercus spp.*) hosts which possess pre-existing gall communities. Natural enemy recruitment is expected to be easier when all of these conditions are met, with the assumption that species are ecologically pre-adapted due to more recently shared evolutionary histories, *and* geographically capable of interacting due to proximity (Schönrogge *et. al..*, 2012). *D. kuriphilus* provides is peculiar since it is distinct in all three of these aspects. Given those traits, we might expect the community of natural enemies to be low or non-existent, yet *D. kuriphilus* is attacked by a rich suite of parasitoids throughout its invaded range, except apparently in the US (Cooper and Rieske, 2007). Nearly all of the parasitoids are idiobiont ectoparasitoids of the superfamily Chalcidoidea, and predominantly members of oak gall communities (see **Table 3** for a summary). The most common *D. kuriphilus* parasitoids in the Palearctic are considered hyper generalists (*Eupelmus urozonus, Torymus flavipes, Bootanomyia dorsalis, Eurytoma brunniventris*)-however all of these comprise sets of morphologically conserved but molecularly distinct lineages of varying host breadths and specialisms, *or* are composed of closely related species-groups that are notoriously difficult to identify (Nicholls *et. al..*, 2017; Gibson and Fusu, 2016; Al-Khatib *et. al.*., 2014). For instance, *B. dorsalis* is composed of two cryptic lineages, one focusing mainly on autumn generation galls, and a second spring specialist which is further sub-divided into ecotypes focusing on *Quercus* section *Cerris* and section *Quercus* hosts respectively (Nicholls *et. al..*, 2017). With a single exception (Ferracini *et. al..*, 2017), molecular identification of *D. kuriphilus* parasitoids has not been implemented in Europe, so we have no idea which cryptic lineages represent *D. kuriphilus*’ natural enemies except for *T. flavipes* (McCormack, 2024), but we should expect the vast proportion of them to be generalists in the stricter sense given the divergence of *D. kuriphilus* from other European oak gallwasps.

Many gallwasp invaders in Europe have been incorporated into native parasitoid host repertoires effectively upon arrival (Schönrogge *et. al..*, 2012), with the exception of asexual galls of *Andricus quercuscalis,* which largely evaded attack for at least twenty years (Hails *et. al..*, 1990). However the effects of establishment time in the increasing diversity of the community, in terms of species richness and rates of attack, are variable. For the most well studied *Andricus quercuscalicis,* there is good evidence of increased species richness and rates of attack by parasitoids (Schönrogge *et. al..*, 1995), but for others there is little data to identify similar trends. Cornell and Hawkins (1993) conducted a meta-analysis on 83 invading herbivorous insects, concluding that, while invaders typically have a lower natural enemy diversity than in their own native range, the cumulative increase in diversity and attack rates with establishment time within the first 150 years of establishment is weakly supported at best.

Parasitoid communities of invading gallwasps may generally resemble their own native communities, excepting the loss of some of their endemic specialists (Schönrogge *et. al..*, 1995). This suggests a degree of non-random ecological filtering processes, imparted upon them by host morphological, phenological and gallwasp host-tree traits (Bailey *et. al..*, 2009). The fact that most if not all of *D. kuriphilus’* native parasitoids are absent from Europe (e.g. Aebi *et. al..*, 2006) means that the European community could not resemble its native community, at least in identity. There is a possibility that similar sets of parasitoids based on their ecologies could assemble, except that currently there are no data of *D. kuriphilus’* native community to assess that. The unique set of trait combinations of *D. kuriphilus* and the extreme divergence in parasitoid species in native and invaded regions make it difficult to predict the structure of its European parasitoid community, although given current data we would expect it to be dominated by hyper-generalists.

### Biological control

A continuing question with respect to *D. kuriphilus* (and to invaders generally) is whether or not native parasitoids are able to impart a regulatory effect. Parasitoids are undoubtedly a major contributor to host mortality in invertebrate herbivore systems (Cornell and Hawkins, 1995), though the extent of population-level control is often not apparent (Hawkins, 1992). For *D. kuriphilus,* some studies report high rates of attack by native parasitoids (e.g. Santi and Maini, 2011), though temporospatial variation is very high, and usually unsatisfactory (Panzavolta *et. al..*, 2013). One of the most promising native species is *T. flavipes,* which remains one of the most common enemies of *D. kuriphilus* (Ferracini *et. al.*., 2018; Matošević and Melika, 2013; Panzavolta *et. al.*., 2013; Santi and Maini, 2011). From currently available data, there is limited support of significant biological control enforced by native parasitoids including *T. flavipes*, although this has only been formally investigated once (Gil-Tapetado *et. al..*, 2021a). Their study showed some indication that after a period of 9 years after establishment of *D. kuriphilus* that there was an increase in rates of parasitoid attack, but the process was disrupted by the massive increase in numbers of the deliberately introduced *Torymus sinensis.* As yet, *T. sinensis* is the only successful form of biocontrol, having been introduced to Japan (Oho and Umeya, 1975), South Korea (Aebi *et. al..*, 2006), Italy (Quacchia *et. al..*, 2008), France (Borowiec *et. al..*, 2018), Spain (Nieves-Aldrey *et. al..*, 2019), Portugal (Amorim *et. al..*, 2022), Croatia (Matošević *et. al..*, 2015), Slovenia (Matošević *et. al..*, 2015), Hungary (Matošević *et. al..*, 2015) and Madeira (Aguiar *et. al..*, 2022). Most recently, introduction trials have also begun in the UK (FERA, 2019). The degree of its success is widespread and dramatic. When *D. kuriphilus* was detected in Italy in 2002, the first European country to be invaded, the chestnut industry around the Piedmont region had effectively collapsed within five years-ten hectares of chestnut orchard declined in value from €300,000 to €30,000 in this time period; equally as dramatically, the industry had recovered almost completely within five years following the introduction of *T. sinensis* (Rondoni, pers comms). Indeed, some of the worst affected areas or northern Italy returned to *D. kuriphilus* population levels so low that they were difficult to detect during surveys (Ferracini, pers. comm.).

Several avenues of biocontrol have been investigated for the reduction of chestnut blight *C. parasitica.* Removal of infected trees or limbs proves ineffective, largely due to the inability to locate all infected trees (EFSA, 2016). Fungicides were equally unsuccessful not only in curbing *C. parasitica,* but also in the non-target effects inherent to their taxonomically indiscriminate nature (Rigling and Prospero, 2018). Chestnut nut rot *Gnomoniopsis castanea,* itself a pathogenic fungus, provides a regulatory effect, but at the unacceptable expense of fruit-damage caused by the fungus itself (Vannini *et. al..*, 2017). The presence of several *Cryphonectria hypovirus* (CHV) species, so-called because of their attenuating effects on *Cryphonectria* have been more successful, though this has only been the case in Europe, where European chestnut is less susceptible to *C. parasitica* than US chestnut (Rigling and Prospero, 2018).

### Non-target effects of *T. sinensis*

Although a very successful biocontrol agent, *T. sinensis* continues to face scrutiny over the degree to which it is a *D. kuriphilus* specialist, as well as to the extent of its hybridization with closely related species. Trials were conducted to test the viability of alternative gallwasp hosts with the conclusion that alternative host utilisation was unlikely (Quacchia *et. al.*., 2008), although these were criticised due to the inappropriate selection of viable species, and for the old age of some of the galls provided (Gibbs *et. al.*., 2011). In a field study, Ferracini *et. al..* (2017) surveyed the surrounding oak gall community where *D. kuriphilus* was present, finding *T. sinensis* emerging from native oak-gall hosts, though at very low levels. Gil-Tapetado *et. al.*. (2023) also find support for non-target ovipositioning into native galls under laboratory conditions, but stressed that only a single individual of *T. sinensis* was reared from native galls in field conditions. Given that *T. sinensis* is so successful at eradicating *D. kuriphilus*, unless *T. sinensis’* populations also collapse and have to be periodically reintroduced during outbreaks, there is a possibility that the absence of its primary host will provide strong selection for the utilisation of alternate hosts. This could effectively lead to a species that may no longer function as an intended biocontrol agent, *and* negatively affect native gallwasp communities. A closely related species, *T. beneficus,* was apparently capable of hybridizing with *T. sinensis,* leading to non-viable biocontrol agents, in addition to concerns as to the disruption of native gallwasp communities with such hybrids (Gibbs *et. al.*., 2011; Aebi *et. al.*., 2006). Further work suggests that *T. sinensis* and *T. beneficus* are composed of three distinct molecular groupings, two of which include *T. beneficus,* and one which includes *T. sinensis* and one of the *T. beneficus* clades (Yara, 2014). In Europe, *Torymus notatus* is another closely related species to *T. sinensis*, whose phenology also overlaps with *T. sinensis* (Pogolotti *et. al..*, 2018), and is also recorded attacking *D. kuriphilus* in several studies (Kos *et. al.*., 2021; Gil-Tapetado *et. al.*., 2021b; Jara-Chiquito *et. al.*., 2019; Ernst, unpublished; McCormack, 2024) hence providing opportunities for hybridization events. So far, there is no evidence of hybridization with native *Torymus* species In Europe (Gil-Tapetado *et. al.*., 2023; Malumphy, pers. comm.), though this may be inevitable with increasing establishment time.

### The purposes of this review

Multiple European countries, and localities within countries have been surveyed for the presence of *D. kuriphilus* and its parasitoid communities. The most comprehensive set of records for the distribution of *D. kuriphilus* were reported by The European and Mediterranean Plant Protection Organisation (EPPO). However, since their primary objectives concerned reporting, preventing or mitigating the spread of *D. kuriphilus*, the level of recording was mostly limited to first recorded establishments at a country or regional level, and they provide no information on its parasitoid community. Within the primary literature of *D. kuriphilus*, many studies have more granular and extensive information that expands beyond the EPPO database. Further, parasitoid community information is scattered across many publications, making it difficult to interpret any broad patterns at a European scale. This chapter incorporates all available data into a unified framework, with additional unpublished data collected by the Food and Environment Research Agency (FERA), Julja Ernst, and from my own UK rearings in 2019 (see Mccormack, 2024). The second objective is to utilise the parasitoid community information to establish any underlying patterns of community assembly as an effect of temporospatial traits. The temporal development of *D. kuriphilus*’ community might suggest that a) native European parasitoids are increasingly competent in utilising a novel resource (*D. kuriphilus*), b) that at some point in the future, native parasitoids may have some regulatory effect on *D. kuriphilus*, and c) implicate the possible negative impact that *D. kuriphilus* may have on native communities via indirect effects such as apparent competition.

Since we already know that *D. kuriphilus*’ main parasitoid enemies are sourced primarily from other oak gall hosts, we should expect that this pattern is reiterated across the entirety of its European range. What is completely unknown is under what governing processes localised communities develop. To date, the assembly of oak gallwasp communities points towards a combination of ecological fitting (similar sets of species assemble based on ecological host traits *i.e* niche assembly; range expansions between parasitoids and their hosts are not strictly bound to each other) and neutral processes: members within a community are a semi-random assemblage governed primarily by rates of migration and extinction events (Bunnefeld *et. al.*., 2018; Stone *et. al.*., 2012). A third scenario is that the couplings between community members are tightly bound, and hence that parasitoids would closely follow the range expansions of their hosts simultaneously, or with some degree of lag (contemporary versus delayed host tracking-Nicholls *et. al.*., 2017). For our purposes, spatial community assembly concerns those processes occurring within the decades that *D. kuriphilus* has established in Europe rather than of evolutionary timescales, and we can assume *a priori* that host pursuit by its native Chinese parasitoids is unlikely to be the main component of community assembly. However, localised host pursuit of *D. kuriphilus* by an initial set of European parasitoids would predict a similar assemblage of species across its European range. If this were true, we should expect components of beta diversity (the difference in composition between local communities) to reflect this. These community-level questions have yet to be explored in *D. kuriphilus*. Our key objectives in the use of the parasitoid data are to answer the following:

1) What are the specific identities of *D. kuriphilus* parasitoids across its European range, and what are their ecological traits?
2) Is *D. kuriphilus* attacked by a consistent set of parasitoids across Europe?
3) To what extent does establishment time bear on the development of its parasitoid communities in terms of species richness and rates of attack?

## Methods

### Literature search

A comprehensive search of all *D. kuriphilus* literature was undertaken using Google Scholar and Web Of Science databases, using the search terms ‘*Dryocosmus kuriphilus’*, ‘chestnut gall wasp’, ‘oriental chestnut gallwasp’ and ‘OCGW’. Since these keywords are certain to appear in any paper concerning *D. kuriphilus*, and because the available literature is relatively limited, every paper was manually examined for relevant content. Each publication’s citations were further examined for hidden sources of information not available from Google Scholar and Web Of Science. Possible grey literature and government websites related to forestry and agriculture and pest management were also investigated using normal Google searches using the same key phrases.

### D. kuriphilus records

The European and Mediterranean Plant Protection Agency (EPPO) served as one of the most comprehensive databases for information on first records and the subsequent spread of *D. kuriphilus* within European countries at regional scales. Additional records were found hidden within primary literature, which sometimes had records at a greater resolution than those reported in EPPO. Many records were reported in multiple sources, and so every record was cross-checked to omit replicated data.

### Parasitoid records

All literature, including any supplemental information were searched for parasitoid records, and recorded within the database in as much detail as possible. Information for parasitoid records were only included if they also provided specific information about the parasitoid identities.

Distribution maps of *D. kuriphilus*, *C. sativa* and parasitoid records were produced using QGIS (v-3.22 ‘Białowieża’), with legend modifications using Inkscape (v-1.1).

### Important terms

The data reported in publications were highly variable. Ideally, all parasitoid community records would be reported at the resolution of a single locality and at a single time point. In reality, studies reported in a range of formats:

1) Multiple localities worth of parasitoid communities were summed together into a single community
2) Multiple years worth of parasitoid data from a single locality were summed together into a single community
3) Combinations of the two

A key term that I define is ‘recording event’. This refers to the highest level of resolution reported for a given community. If a paper only reported a single community summed across a range of years (i.e. abundance data from a single location is the sum of multiple years of data), then this is reported as a single recording event. Corresponding information, and further definitions that capture the nature of the data are reported in **Appendix 1**. For data modelling purposes, the resolution of the data was taken into account and filtered appropriately (described below).

### Statistics- Measuring spatiotemporal components of diversity

The diversity of communities are generally considered at 3 levels of resolution: alpha, beta and gamma diversity. Alpha diversity concerns the diversity of a community at a local scale. This could be a community within a single locality, or to a community measured at a single time point. Gamma diversity concerns the total diversity of a set of local communities. This is the diversity captured across the entire set of sampled localities (i.e. regional scales), the entire set of sampled time points of a locality, or a combination of the two. Beta diversity concerns the differences of diversity between local communities, and by extension, the relationship between local diversity (alpha) and total diversity (gamma). All components of diversity are further delineated based upon their measurements of diversity:

1) Zeroth-order diversity considers only the presence or absence of members of a given community. This is the raw number of species within a given community at an alpha and gamma level, or the difference between the membership of species between different communities (*i.e.* beta-diversity dissimilarity).
2) First-order diversity is the membership of communities with an additional quantification of its members, most commonly by their abundance or biomass.

### Alpha diversity

For this study, core components of alpha diversity that are implemented for statistical purposes are the exponent of the Shannon entropy, while for descriptive purposes the raw abundance or presence (when abundance data were not available) are used. The Shannon entropy (H) consists of the following formula:

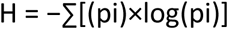

Where H is the Shannon index, and pi refers to the proportion (p) of individuals of a given species (i) relative to the total number of individuals of all species of a community (ni/N). The Shannon entropy in an untransformed state is an index of diversity, which considers the richness of a community (the total number of species) in addition to a measure of their evenness (whether all species are equally abundant or not). However, the values produced are difficult to interpret on their own and to compare with values from other communities. The Shannon entropy is commonly transformed using the exponent of the resulting value to create the ‘effective number of species’ (Tuomisto, 2010a). This value provides an intuitive sense of the number of species that are equally likely to be sampled at random. The value is bound between 1 (assuming that at least 1 species has been sampled for a given community) and the total number of species sampled in a community. For illustration, consider a community sample where there are 10 species in total, but 9 of those species were only represented by a single individual, and 1 species was sampled 10,000 times-then the effective number of species would be approximately equal to 1. Conversely, if we sampled another community with 10 species, and 100 individuals of each of those 10 species, then the effective number of species would be exactly 10.

### Beta Diversity

Traditional beta-diversity by Whitaker was to consider it as the ratio between *mean* alpha diversity (the average diversity of local communities) and gamma diversity (the combined diversity of all communities-Tuomisto, 2010b), or the ‘effective number of communities’: beta = gamma / mean(alpha). Local communities that are identical to the larger community as a whole, have no compositional difference between them and therefore 0 beta-diversity (Tuomisto, 2010b). In such a case, alpha diversity is equal to gamma diversity. Another way to consider the effective number of communities is to think of the value as an answer to the question: how many local communities would I need to randomly sample in order to fully sample all the members of the total community? Classical beta-diversity indices include the Sörensen or Jaccard dissimilarity index, which are zeroth order metrics that focus on the presence or absence of species, and the first order Bray-Curtis dissimilarity index which is abundance-based (Baselga 2017; Baselga and Orme, 2012; Tuomisto, 2010b). All dissimilarity indices are bound between 0 and 1, with 0 indicating communities with no compositional difference between them, and 1 being complete dissimilarity. From here on I use the Sørensen index. The Sørensen index has a greater emphasis on the species that *are* shared rather than to the total number of species that are not.

The Sørensen index is divided into two distinct components by which communities can differ:

1) A set of communities typified by the strict addition or loss of species-in other words, the species within the poorest community are a strict subset of the richest i.e. the poorest community has no species that are unique compared to other communities (nestedness-resultant dissimilarity).
2) A set of communities dominated by the replacement of species for others i.e. communities all have some number of species that are completely unique (turnover-resultant variation).

Inherent within a nested community is the notion that local communities *must* have a difference in species richness. This is not a necessary component of a community dominated by turnover however. For example, 3 local communities could share 3 species in common, and each have 3 species unique to each, and therefore each have 6 species in total (hence no richness difference).

The inclusion of species abundance in a Bray-Curtis index adds 2 additional metrics of community difference:

1) Communities where the total abundance of each community is identical, but where there are differences in the relative abundance of each species (balanced variation).
2) Communities where the relative abundance of each species is uniform across local units, but where the total abundance of local communities vary (gradient-resultant variation).

A community that differs in balanced variation might suggest something about differing dominance structures for example, whereas gradient based variation might say something about the total availability of resources along an ecological cline. In reality, communities usually display a combination of all of the above components, and modern metrics allow some quantification of these related but distinct qualities to provide more informative ways of interpreting community difference.

### Computing beta-diversity

In order to compute the different components of compositional change in *D. kuriphilus* (Sørensen, turnover, nestedness, Bray-Curtis, balanced, gradient), the functions beta.pair() and beta.pair.abund() were used from the betapart package (v1.6-Baselga and Orme, 2012) in R platform (v-4.3.2).

### Testing the consistency of community composition-multiple regression on matrices (MRM)

Multiple regression on distance matrices (MRM) were developed in order to allow inferential statistics of distance matrices in relation to environmental and trait influences in much the same way that any other multiple regression works (Lichstein, 2007). In common with multiple linear regressions, MRMs directly adopt a regression approach with different permutation methods to estimate the statistical significance of the parameter estimates (Lichstein, 2007). Inherent within studies investigating community change with geographic distance for example, is the possibility of differences in the magnitude of dissimilarity at different spatial scales. Lichstein (2007) adopts the use of lag matrices: the grouping of geographic pairwise distances into distinct distance classes. The inclusion of these lags into the model allows for non-linearity in the data, by applying a regression to each lag separately, and comparing them to the mean of all lags. Lags that are significantly less than or greater than the mean provide evidence of a stronger effect at different spatial scales than would otherwise be expected with a purely linear relationship. Statistical significance is adjusted using a Bonferroni correction to account for multiple comparisons. The appropriate number of lags is calculated using Sturge’s rule based on the total number of elements within the matrix (Lichstein, 2007). Lag divisions are quantile-based (the size of lags are determined by partitioning an approximately equal number of pairwise comparisons within each lag, i.e. the distance between lags can vary but the number of pairwise comparisons cannot), or the entire geographic distance is split into equally sized distance classes (i.e. the number of pairwise distances within each lag can vary but the lengths of the lags cannot). For this review, quantile-based lags are used. Mantel correlograms were performed on each of the lags and compared to the mean of all lags, a separate coefficient for each lag was estimated and then tested for significant differences (**Figure 2**). The functions ‘mgram’ and ‘MRM’ in the R package ecodist (v 2.1.3-Goslee and Urban, 2007) were used to perform Mantel correlograms and MRM analyses respectively. For the results, only models produced from data where the number of galls and parasitoid abundance were reported are included (‘DK3’ dataset).

**Figure 2.**
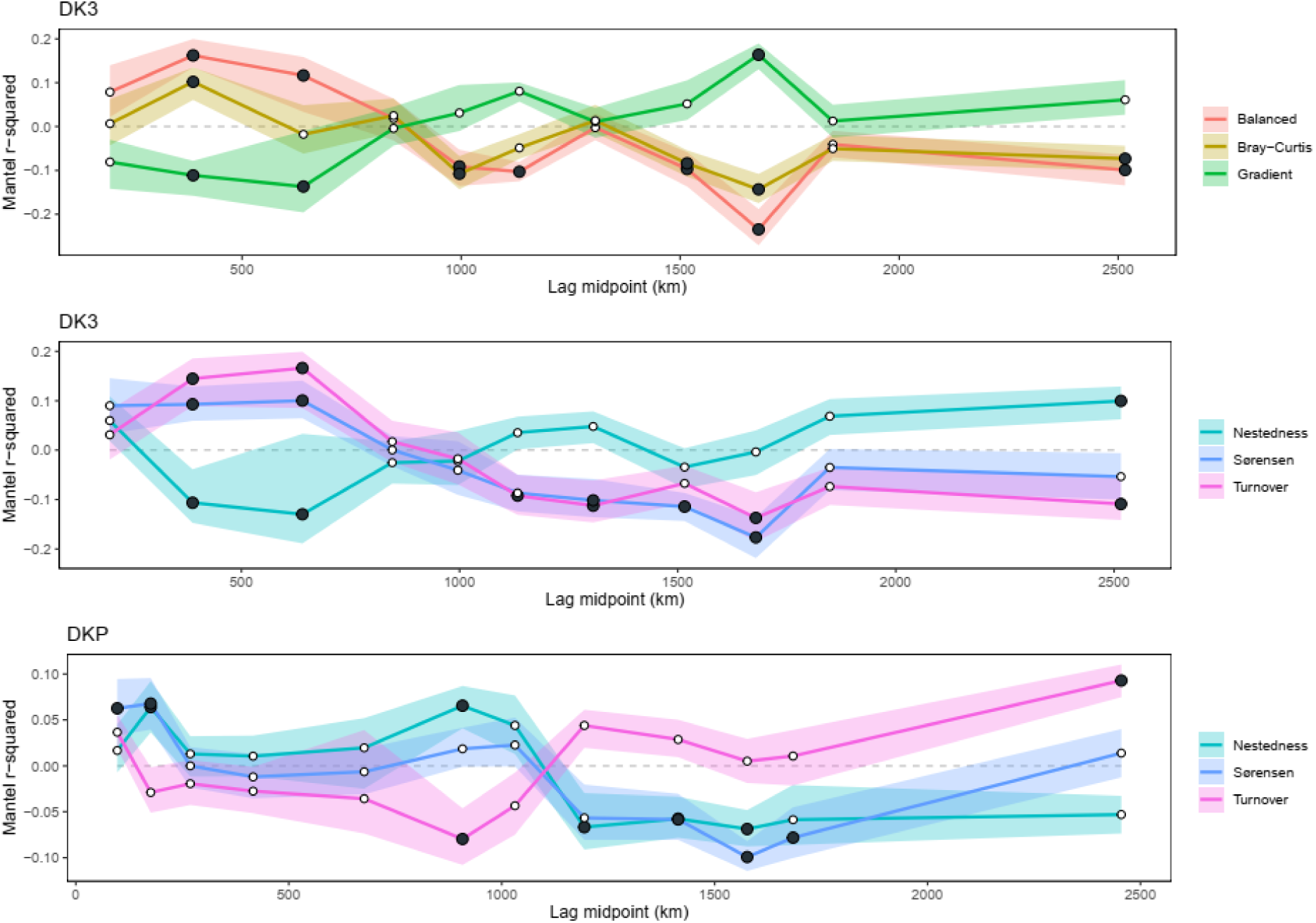
Lag point Mantel correlograms of significance for beta diversity metrics in order of major data structure. Lag distances are split into quantiles with an equal number of pairwise distances within them, and hence, the distances between 1 lag and another is allowed to vary. Above and below each dissimilarity component are coloured ribbons representing the upper and lower confidence intervals about each lag estimate coefficient. ‘DK3’ data structure includes only communities where the number of galls *and* the number of parasitoids were reported. The uppermost graph displays abundance based beta diversity metrics, whereas the central graph displays presence absence beta diversity metrics. DKP data structure includes additional communities where the number of galls reared were reported but not community abundance, plus all communities from DK3. Black circles indicate lags which are statistically significantly different to the mean correlation coefficient of all lags, and white circles are non-significantly different.

For each respective component of beta-diversity, only distance lags that were statistically significantly different to the total lag mean (results in **Figure 2**) were included as predictors in their final models. In all models, the difference between the log number of galls reared, and the elapsed time of establishment are included as additional explanatory terms of community dissimilarity metrics.

For the production of geographic distance matrices, the st_as_sf() function was used in the sf package (v1.0-15: Pebesma, 2018) within R to convert raw latitude longitude coordinates into geodesic distances.

### Data criteria and cleaning

Two differently sized datasets based on differing parameters for beta diversity analyses were used. All analyses had common omissions. Community data were only included if all species were reported. For instance, Kos *et. al..* (2020) only reported the 7 most abundant parasitoids reared. Ferracini *et. al..* (2017) fully described the community, but reduced all individuals of *Eupelmus* into *Eupelmus sp,* on the basis that *Eupelmus* is composed of many morphologically conserved but molecularly distinct, or very difficult to identify species. Although this is true, morphospecies information would have made it easier to directly compare with other papers, and narrowed the number of possible cryptic lineages that any *Eupelmus sp.* could have been. For example, *E. urozonus* is a known and very common parasitoid of oak gallwasps with numerous cryptic lineages (Ernst, unpublished; Al-Khatib *et. al.*, 2014; Quacchia *et. al.*, 2012). By knowing that individuals of *Eupelmus* belong to the *E. urozonus* complex, we can at least know that its primary hosts are oak gallwasps rather than some species otherwise commonly reared from non-oak-gall hosts. Ferracini *et. al.* (2017) was otherwise a study rich in community information (34 communities) comprising a significant proportion of the total available communities for analysis. Numerous ways of dealing with the data were considered:

1) To ignore all the communities from Ferracini *et. al.* (2017).
2) Reduce all species of *Eupelmus* from all papers to *Eupelmus sp*.
3) Assign *Eupelmus* sp within the Ferracini *et. al.* (2017) paper to the most likely species which is *E. urozonus,* or portion them out as a proportion of the likely species based on other communities.
4) Acknowledge the problem and carry on with the analysis regardless.

I used Mantel correlation tests (Goslee, 2010) within the ‘ecodist’ package (v-2.1.3-Goslee and Urban, 2007) in R, which are a Pearson’s correlation-equivalent for multivariate data, to estimate the amount of change in dissimilarity scores between full species dataset and the same dataset where *Eupelmus spp* are reduced to *Eupelmus sp* (**Table 1**). The conclusion was that the data were relatively robust and that any choice made was unlikely to alter the overall conclusions made in subsequent analyses. The final analyses implement data where all *Eupelmus spp.* were transformed into a single ‘*Eupelmus sp.’* taxon.

**Table 1.**
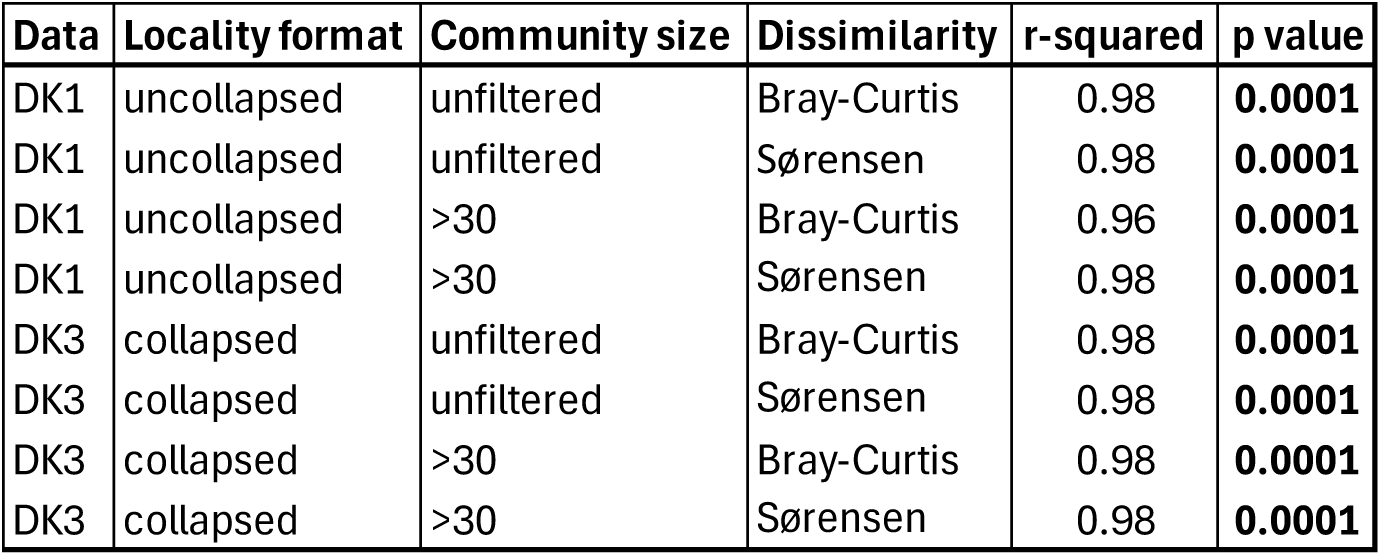
Mantel correlation tests between all species vs *Eupelmus-*reduced equivalents, and under a series of different data-manipulations. ‘Locality format’ relates to whether or not repeat localities were collapsed into single communities or not (DK1 and DK3 ‘Data’ respectively). ‘Community size’ relates to whether or not communities with less than 30 individuals were omitted or not. ‘Distance measure’ relates to the measure of dissimilarity (Sørensen or Bray-Curtis). Bray-Curtis is abundance-based and Sørensen is presence-absence based.

### Testing for community change through time

Mixed effects models were used to explore the possible relationship of increased parasitoid abundance, and the effective number of species as an effect of the length of establishment of *D. kuriphilus*.

*D. kuriphilus* may incur increased rates of attack with elapsed establishment time, as an adaptive response from the surrounding community of available parasitoids to exploit a novel resource (Cornell and Hawkins, 1993). If this were true, we should expect the fraction of parasitised versus unparasitised *D. kuriphilus* galls to increase over time. In order to answer this we would need some estimate of parasitised versus unparasitised galls in addition to a measure of host population size, but the most commonly available data available were the number of galls reared, versus the number of parasitoids reared. While these are not ideal metrics, our expectation should be that these two measures would broadly reflect the underlying pattern of increasing parasitoids per unit gall per unit time. Hence the inclusion of the number of galls reared, along with the years since *D. kuriphilus* was first recorded within a given region are the primary predictors of parasitoid abundance.

There may be many other factors influencing parasitoid abundance such as climatic (Gil-Tapetado *et. al..*, 2021a; 2020; Bonsignore *et. al..*, 2020) or geographic factors. Within the European range of *D. kuriphilus* are numerous regions which were putative glacial refugia from the Last Glacial Maximum (Habel *et. al.*., 2010), which generally have a higher diversity of species than surrounding areas. The availability of climatic data or otherwise is limited, but the hope was that some of these effects would be captured by the region as levels in a predictor. However, the level of replication for regions across different time points is too sparse to reliably estimate regional effects. Further, the variability of different regions is large. In order to account for this, a linear mixed regression was fitted with ‘Region’ as a random effect, and the number of galls reared and time since establishment as fixed effects.

*D. kuriphilus* may have a richer community of common parasitoid species as a function of establishment time elapsed, which could be due to adaptive processes from the parasitoids’ utility of a novel resource, as a consequence of increased detection rates due to *D. kuriphilus’* increasing population size, stochastic detection processes, or all of the above (Cornell and Hawkins, 1993). The parasitoid abundance regression makes no assumptions about the membership of the parasitoids. A single parasitoid species could be responsible for the increased rates of attack, *or,* the increased rates of attack could be the result of additive rates of attack from a set of parasitoid species. In the latter case, the effective number of parasitoid species should increase as a consequence of time of establishment. As with the parasitoid abundance regression, a linear mixed model was fitted with Region as a random effect, number of galls reared and years since first record as fixed effects.

Linear mixed effects were performed using the lmer() function within the R package lme4 (v 1.1-Bates, 2010), and significance tests were computed using an expansion of the base R function summary() using the package lmerTest (v 3.1-Kuznetsova *et. al.*., 2017). Model checking of residuals and overdispersion were performed using the plot() function in base R, and testDispersion() within the R package DHARMa (v. 0.4.6-Hartig and Hartig, 2017).

## Results

From 2002-2020, a minimum of 1,017,060 galls were reared across Europe from 32 publications and 3 unpublished studies (**Appendix 2**) comprised of 178 localities across 11 countries (**Fig 3**; **Table 2**). Six studies did not report on the number of galls reared, and 3 provided partial reports (Gil-Tapetado *et. al..*, 2021b; Bonsignore *et. al.*., 2019; Bonsignore and Bernardo, 2018; Bernardinelli *et. al.*., 2016; Boriani *et. al.*., 2013; Speranza *et. al.*., 2008; Aebi *et. al..*, 2006; Ernst, unpublished; Malumphy, unpublished), hence the true number of galls reared is greater than the value reported here. Over half of communities sampled were in Italy (93 communities), and just over a third of the remaining from Spain (41 communities) and England (21 communities). 67% of all studies included community data spanning a single year. The next 20% of studies included repeated sampling of 2 and 3 years, and the remaining with 4, 5 and 6 years of data. Detailed probability distribution estimates of *C. sativa* indicate very strongly that *D. kuriphilus* occupies many regions with chestnut presence probability estimates above 5% (**Figure 3 (below)**).

**Fig 3.**
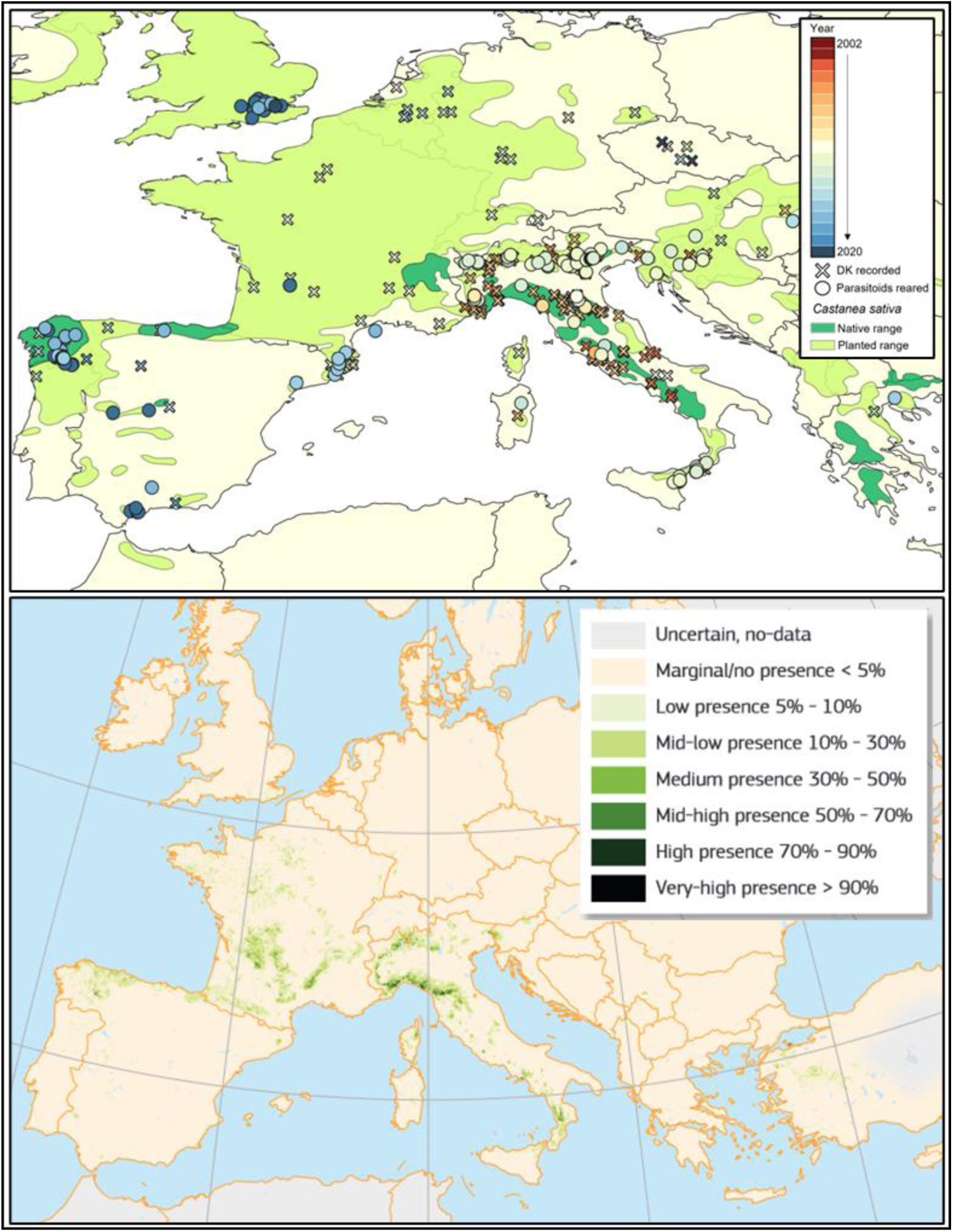
The distribution of *D. kuriphilus,* parasitoid records and *C. sativa* distribution (above), and probability estimates of *C. sativa* presence in Europe (below). *C. sativa* shape files in the upper map were taken from Caudullo *et. al..* (2017). *D. kuriphilus* records are coloured by year of detection, or by date of rearing of parasitoids (cross and circle symbols respectively). The lower map is modified from Conedara *et. al..* (2016).

**Table 2.** Community sampling across Europe. Countries are ordered in descending order of the total number of recording events (see **Methods-Important terms**). “No.” refers to the number of recording events, with a corresponding percentage of total recording events for a given country and year.

|  | Years of repeated data collection |  |  |  |  |  |  |  |  |  |  |  |  |  |
| --- | --- | --- | --- | --- | --- | --- | --- | --- | --- | --- | --- | --- | --- | --- |
| Country | 1 |  | 2 |  | 3 |  | 4 |  | 5 |  | 6 |  | Total |  |
|  | No. | % | No. | % | No. | % | No. | % | No. | % | No. | % | No. | % |
| Italy | 74 | 41.6 | 12 | 6.7 | 4 | 2.25 | 1 | 0.56 | 1 | 0.56 | 1 | 0.56 | 93 | 52.2 |
| Spain | 17 | 9.6 | 1 | 0.6 | 15 | 8.43 | 8 | 4.49 |  | - |  | - | 41 | 23.0 |
| England | 20 | 11.2 |  | - |  | - | 1 | 0.56 |  | - |  | - | 21 | 11.8 |
| Croatia |  | - | 4 | 2.2 | 1 | 0.56 |  | - |  | - |  | - | 5 | 2.8 |
| Slovenia | 2 | 1.1 |  | - | 1 | 0.56 | 2 | 1.12 |  | - |  | - | 5 | 2.8 |
| Turkey | 5 | 2.8 |  | - |  | - |  | - |  | - |  | - | 5 | 2.8 |
| France |  | - |  | - |  | - | 1 | 0.56 | 1 | 0.56 |  | - | 2 | 1.1 |
| Portugal | 1 | 0.6 |  | - |  | - | 1 | 0.56 |  | - |  | - | 2 | 1.1 |
| Switzerland |  | - | 1 | 0.6 |  | - |  | - | 1 | 0.56 |  | - | 2 | 1.1 |
| Greece | 1 | 0.6 |  | - |  | - |  | - |  | - |  | - | 1 | 0.6 |
| Hungary |  | - |  | - | 1 | 0.56 |  | - |  | - |  | - | 1 | 0.6 |
| Total | 120 | 67.4 | 18 | 10.1 | 22 | 12.36 | 14 | 7.87 | 3 | 1.69 | 1 | 0.56 | 178 | 100.0 |

### Parasitoid communities

Of the studies that included abundance information, 132,352 parasitoids were recorded. 87,367 of these were *T. sinensis*, and 44,985 were native parasitoids (**Table 3**). A total of 88 taxa have been recorded attacking *D. kuriphilus* across the European range, with 72 confirmed to species-level. Taking into account only presence-absence, and with the inclusion of all taxa including those only confirmed to a generic level, the *D. kuriphilus* community is overwhelmingly represented by parasitoid species attacking galling insects (87.4%), of which the vast majority are known from oak gallwasp hosts (70.1% oak gallwasps, 17.2% non-oak galls). Including only taxa confirmed to species level, 90.4% are gall specialists, of which 82.2% are oak gall specialists. The remaining parasitoid taxa are comprised of specialists of leaf miners, borers, solitary hymenopterans, eggs and unknown insect hosts. Excluding *T. sinensis* and taking account of abundance, just over 99.7% of parasitoids reared are gall parasitoids, 99.2% of which are oak gall specialists.

**Table 3.**
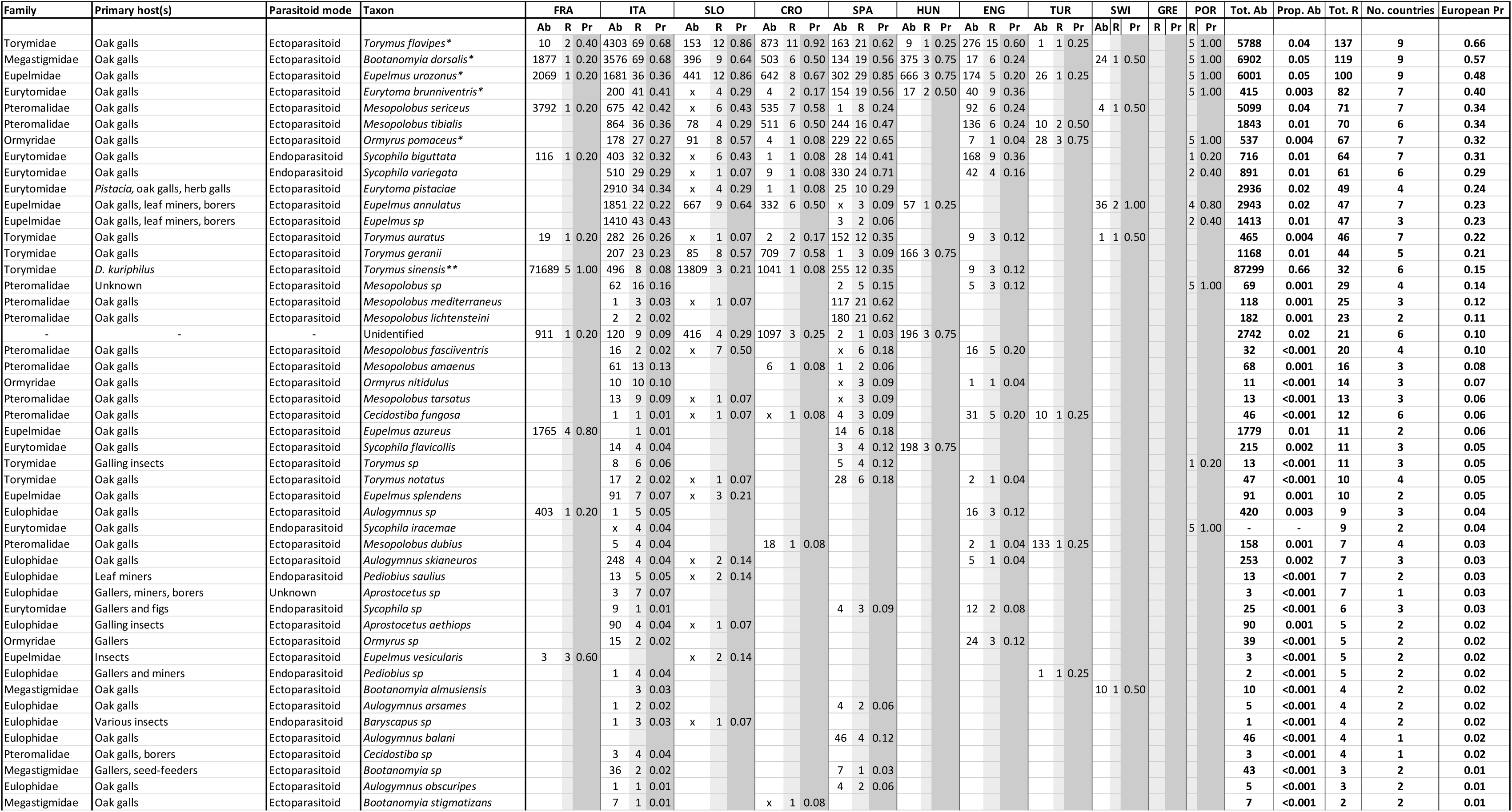
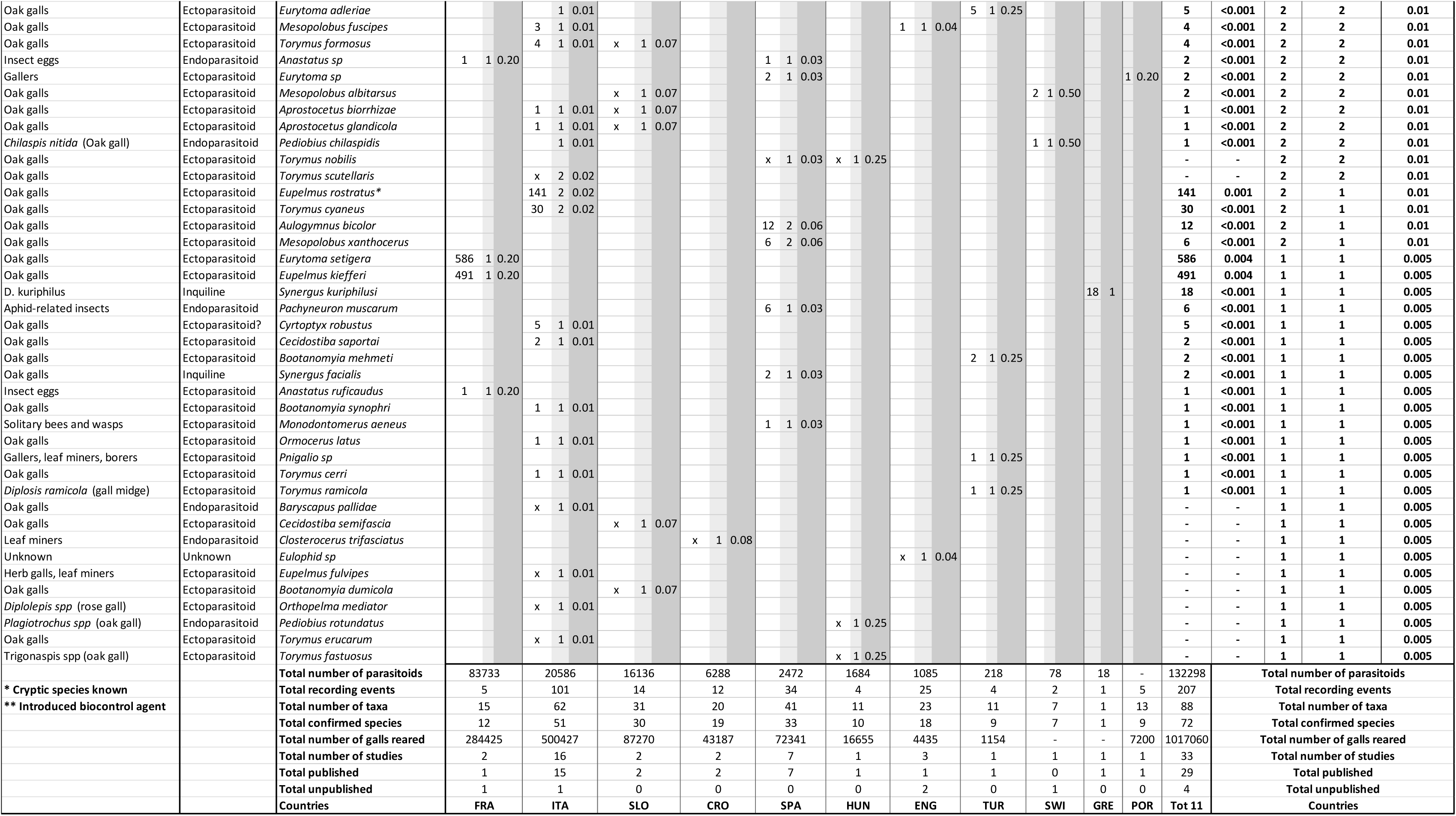
The parasitoids of *D. kuriphilus* within its European range. Country columns are ordered in descending order of total insect abundance from left to right. Rows are ordered in descending order of European prevalence (‘**European Pr’** = Total country recording events of a parasitoid / Total European recording events. See **Methods-Important terms**). Country codes are as follows: **FRA** = France, **ITA** = Italy, **SLO** = Slovenia, **CRO** = Croatia, **SPA** = Spain, **HUN** = Hungary, **ENG** = England, **TUR** = Turkey, **SWI** = Switzerland, **GRE** = Greece, **POR** = Portugal. **Ab** = Abundance, **R** = number of Records, **Pr** = Number of records of a species in a country / Total number of country recording events. **Prop. Ab** = Total species abundance/ Total European abundance. **No. countries** = Number of countries with records of a given species, **European Pr** = European Prevalence = Total number of species recording events / Total number of European recording events.

Most parasitoid species belong to the superfamily Chalcidoidea (96.6%), with the remaining Cynipoidea (*Synergus* inquilines-1.2%), Ichneumonoidea (1.2%) and unidentified (1.2%). Parasitoid representatives are predominantly ectoparasitoids (79.1%), followed by endoparasitoids (15.2%), and the remaining unknown or inquilines. While ectoparasitoid koinobiont species do exist, all species recorded here are idiobionts, although Askew (1975) states that some species of *Aulogymnus* and *Torymus cyaneus* (both recorded attacking *D. kuriphilus*) completely avoid paralysing their hosts, living ectoparasitically with the still developing host larva, echoing some characteristics of koinobiosis.

The most common native parasitoid species attacking *D. kuriphilus* are *T. flavipes*, *B. dorsalis, E. urozonus, E. brunniventris, M. sericeus* and *M. tibialis* (**Table 3**, **Figure. 5**), all amongst the most common parasitoids across the Western Palearctic (Askew *et. al.*, 2016).

### Community consistency

Across Europe, the community of *D. kuriphilus* is highly variable. The relationship between beta-dissimilarity and distance is non-linear, with total dissimilarity (Bray-Curtis and Sørensen dissimilarity) reaching an asymptote of maximum dissimilarity between 900km and 1200km (**Figure 4**). Contrastingly, the relative contributions of turnover and balanced-resultant dissimilarity components, compared to nestedness and gradient resultant dissimilarity, increase noticeably with increasing geographic distance.

**Figure 4.**
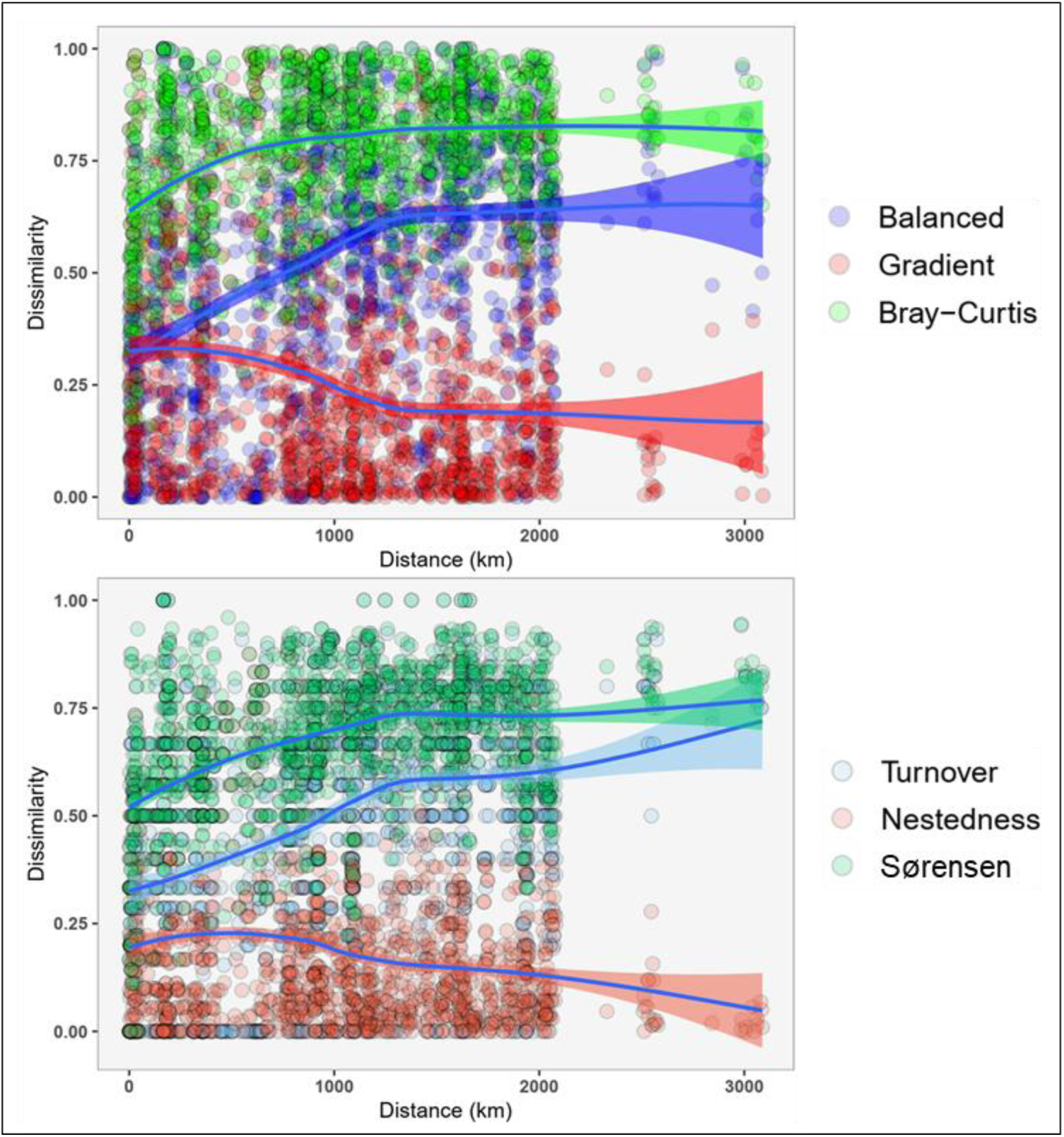
Components of abundance-based (above) and presence-absence based (below) beta diversity in relation to geographic distance. Results are reported for communities where the number of galls reared and parasitoid abundances were recorded (N communities = 55, pairwise comparisons = 1485). Locally estimated scatterplot smoothing (LOESS) is fitted to indicate the non-linear relationship of beta diversity components with increasing geographic distance.

### Statistical analyses- Multiple regression on distance matrices (MRM)

Each MRM included distance lags with significant spatial autocorrelation from the results of the Mantel correlograms for each respective dissimilarity component (**Methods- Figure 2**), plus the number of galls reared (log transformed) and the years since first record that *D. kuriphilus* was detected as additional predictors (**Table 4**). The percentage of variance explained by the models was generally low (≤ 18%), though highest when accounting for parasitoid abundance (i.e. Bray-Curtis total dissimilarity and its components: balanced and gradient-resultant dissimilarity). Balanced-resultant dissimilarity increased significantly at lag distances with midpoints of 996.1km (Lag6), and from 1515.8km, 1679.1km and 2516km (Lags 9, 10 and 12). Neither the number of galls reared nor the years since first record of *D. kuriphilus* had a statistically significant impact on balanced-resultant dissimilarity when accounting for lag classes. Without the consideration of parasitoid abundance, turnover-resultant dissimilarity explains more than half of the variance explained that balanced-resultant dissimilarity does (10% vs 17%). Similarly, turnover increases with increasing geographic distance, significantly from 1133km (Lag7), 1306.3km (Lag8), 1679.1km (Lag10) and 2516km (Lag12). Gradient-resultant dissimilarity increases with the number of galls reared, and years since first record of *D. kuriphilus*.

**Table 4.**
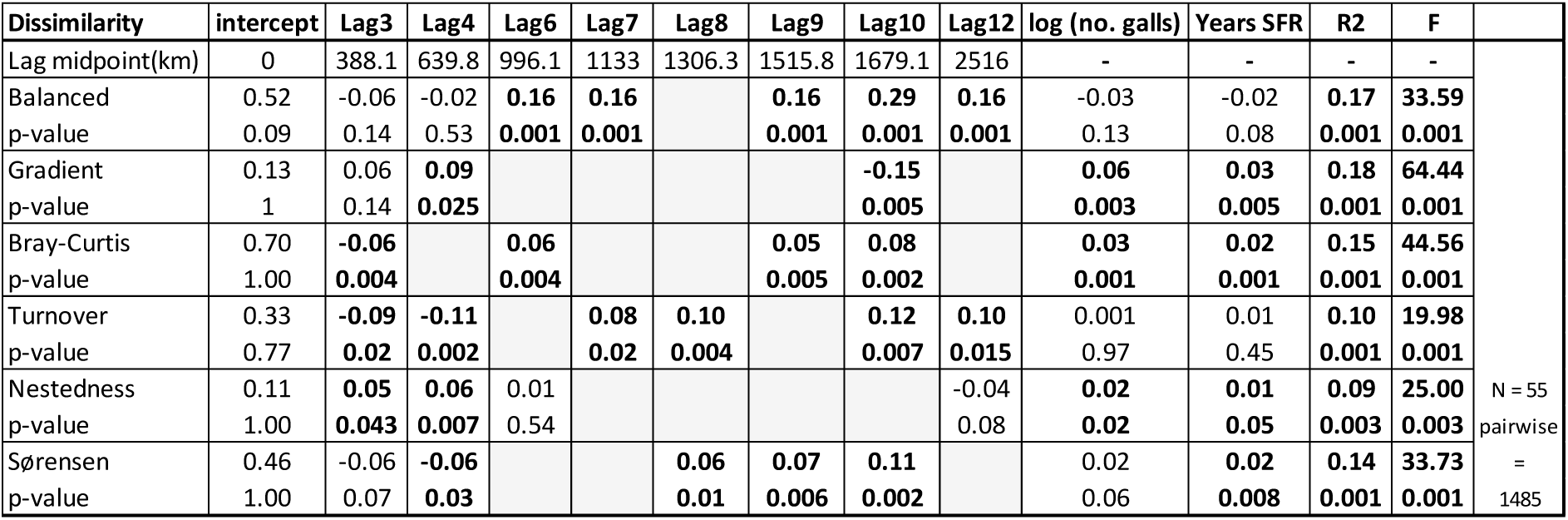
Multiple regression on distance matrices for beta diversity components. Some lags are omitted because they showed no statistically significant spatial autocorrelation for any components of beta diversity. Years SFR = years since first record. Significant coefficients are indicated in bold, with their respective p-values reported immediately below them. Lags not included for a given beta-diversity component model are left blank and coloured in grey.

| Dissimilarity | intercept | Lag3 | Lag4 | Lag6 | Lag7 | Lag8 | Lag9 | Lag10 | Lag12 | log (no. galls) | Years SFR | R2 | F |  |
| --- | --- | --- | --- | --- | --- | --- | --- | --- | --- | --- | --- | --- | --- | --- |
| Lag midpoint(km) | 0 | 388.1 | 639.8 | 996.1 | 1133 | 1306.3 | 1515.8 | 1679.1 | 2516 | - | - | - | - |  |
| Balanced | 0.52 | -0.06 | -0.02 | **0.16** | **0.16** |  | **0.16** | **0.29** | **0.16** | -0.03 | -0.02 | **0.17** | **33.59** |  |
| p-value | 0.09 | 0.14 | 0.53 | **0.001** | **0.001** |  | **0.001** | **0.001** | **0.001** | 0.13 | 0.08 | **0.001** | **0.001** |  |
| Gradient | 0.13 | 0.06 | **0.09** |  |  |  |  | -0.15 |  | **0.06** | **0.03** | **0.18** | **64.44** |  |
| p-value | 1 | 0.14 | **0.025** |  |  |  |  | **0.005** |  | **0.003** | **0.005** | **0.001** | **0.001** |  |
| Bray-Curtis | 0.70 | -0.06 |  | **0.06** |  |  | **0.05** | **0.08** |  | **0.03** | **0.02** | **0.15** | **44.56** |  |
| p-value | 1.00 | **0.004** |  | **0.004** |  |  | **0.005** | **0.002** |  | **0.001** | **0.001** | **0.001** | **0.001** |  |
| Turnover | 0.33 | -0.09 | -0.11 |  | **0.08** | **0.10** |  | **0.12** | **0.10** | 0.001 | 0.01 | **0.10** | **19.98** |  |
| p-value | 0.77 | **0.02** | **0.002** |  | **0.02** | **0.004** |  | **0.007** | **0.015** | 0.97 | 0.45 | **0.001** | **0.001** |  |
| Nestedness | 0.11 | **0.05** | **0.06** | 0.01 |  |  |  |  | -0.04 | **0.02** | **0.01** | **0.09** | **25.00** | N = 55 |
| p-value | 1.00 | **0.043** | **0.007** | 0.54 |  |  |  |  | 0.08 | **0.02** | **0.05** | **0.003** | **0.003** | pairwise |
| Sørensen | 0.46 | -0.06 | -0.06 |  |  | **0.06** | **0.07** | **0.11** |  | 0.02 | **0.02** | **0.14** | **33.73** | = |
| p-value | 1.00 | 0.07 | **0.03** |  |  | **0.01** | **0.006** | **0.002** |  | 0.06 | **0.008** | **0.001** | **0.001** | 1485 |

Gradient-resultant dissimilarity has similar explanatory power to Balanced-resultant dissimilarity (18% vs 17%), though its base estimate (0.13, p = 1) indicates that its relative contribution to dissimilarity is not significantly different to 0 on average. Its highest contribution to dissimilarity occurs at a lag midpoint of 639.8km (Lag4), and dramatically decreases at 1679.1km (Lag10) to an effective contribution to community dissimilarity of 0 (**Lag10 = −0.15, p = 0.005**). Both the number of galls (**log (no. galls) = 0.06, p = 0.003**) and years since first record (**years SFR = 0.03, p = 0.005**) increase the contributions of gradient-resultant dissimilarity, suggesting that similar sets of species are more likely to be sampled with increasing sample size and time of establishment. Nestedness-resultant dissimilarity bears a similarly minimal contribution to overall dissimilarity, and is also significantly increased by a small amount with increasing sample size (**log (no. galls) = 0.02, p = 0.02**) and length of establishment (**Years SFR = 0.01, p = 0.05**).

The most commonly encountered parasitoids across communities are *T. flavipes, B. dorsalis* and *Eupelmus sp* (**Figure 5**), of which the majority of *Eupelmus sp* are represented by the morphospecies *E. urozonus* (**Appendix 3**). The dominance structure of the three most common parasitoids varies considerably, although *T. flavipes* is most commonly the dominant species in Europe. Spanish communities are noticeably higher in the number of parasitoids attacking *D. kuriphilus*, and most divergent compared to other European communities. Of important mention however is the existence of *T. sinensis* in Spanish communities, which dramatically influences the structure of communities where it is prevalent (Gil-Tapetado *et. al.*., 2021a).

**Figure 5.**
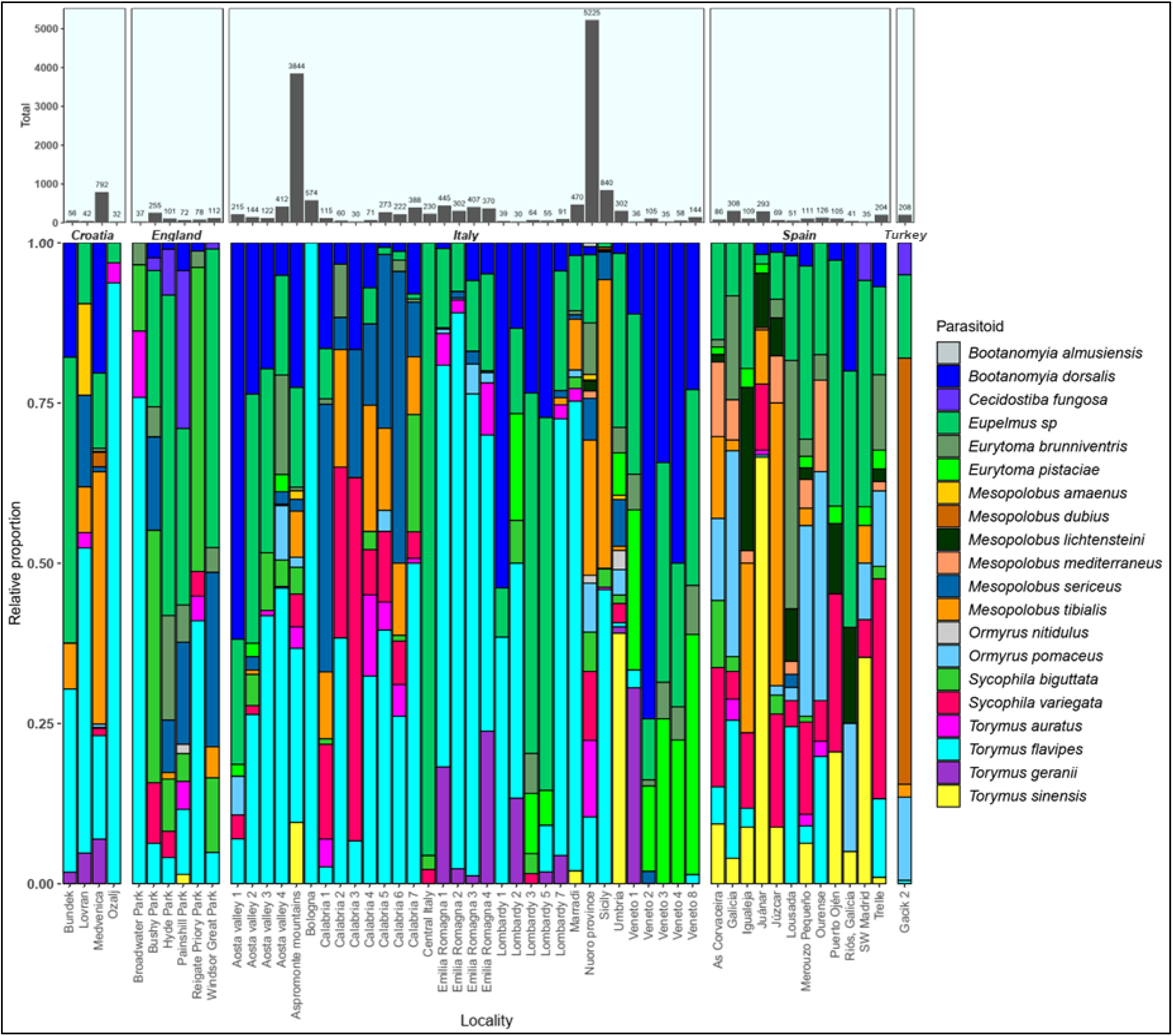
The relative proportion of the top 20 parasitoid species of *D. kuriphilus* across Europe (below) and the number of parasitoids reared (top). Localities where less than 150 galls were reared have been excluded for brevity. The total number of parasitoids reared are labelled above each corresponding bar. The data included are from the dataset where Ferracini *et. al.*. (2017) communities are included, and *Eupelmus spp* are reduced to *Eupelmus sp.* See also **Appendix 3** for communities that include all *Eupelmus spp* information.

### Abundance as an effect of length of establishment

After accounting for the variance contained within geographic regions (**Table 5-Random effects**), and with the inclusion of sample size and time of establishment as fixed effects, the only statistically significant influence on parasitoid abundance was sample size (**log (No. galls): Estimate = 2.02, p = <0.0001**). Time of establishment has an effect of increased abundance of 0.92 parasitoids with each increasing year, but this is not statistically significant (Years since first record: p = 0.27).

**Table 5.** Mixed effects model of abundance as an effect of time of establishment. Coefficient values are exponents of their original values because the model was fitted to log abundance.

| Random effects: |  | Variance | Std.Dev. |  |  |
| --- | --- | --- | --- | --- | --- |
| Region | (Intercept) | 0.93 | 0.97 |  |  |
| Residual |  | 1.62 | 1.27 |  |  |
| N obs, N groups | 79,7 |  |  |  |  |
| Fixed effects: | Estimate | Std. Error | df | t | p-value |
| (Intercept) | 0.76 | 0.98 | 51.40 | -0.29 | 0.78 |
| log(No. galls) | 2.02 | 0.14 | 75.94 | 5.03 | <0.0001 |
| Years since first record | 0.92 | 0.07 | 72.67 | -1.11 | 0.27 |

### Effective number of species as an effect of length of establishment

After accounting for the variance within geographic regions (**Table 6-Random effects**), and sample size and time of establishment as fixed effects, the only statistically significant influence on the effective number of species was sample size (**log (No. galls): Estimate = 0.85, p = <0.0001**). The effective number of species increases by 0.12 species with each increasing year of establishment, but this was not statistically significant (Years since first record: p = 0.18).

**Table 6.**
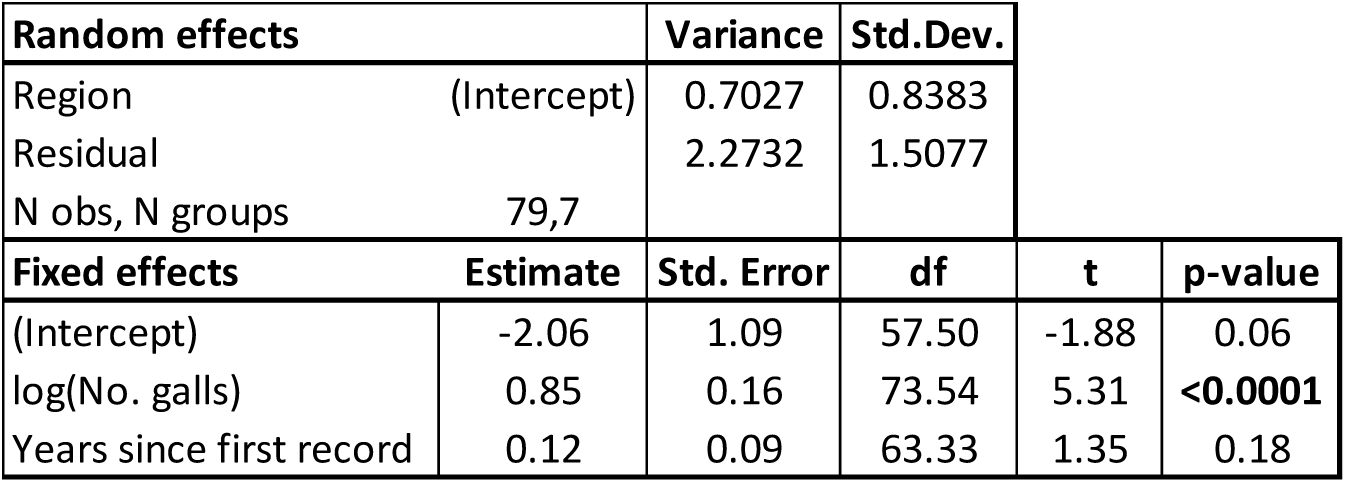
Main effects of the final effective number of species model (ANOVA) and associated values from a multiple regression.

## Discussion

### The distribution of *D. kuriphilus*

Based upon the distribution of *C. sativa* within Europe, *D. kuriphilus* has expanded into most regions bearing significant stands of sweet chestnut within less than 3 decades. Survey efforts of *D. kuriphilus* have been extensive, although the primary focus on *D. kuriphilus* as an economic pest has led to the most extensive surveys surrounding areas of significant chestnut industry and little elsewhere. The expanse of records surrounding these areas, particularly in Italy, France, Spain and Portugal are apparent. The extent of *C. sativa* (**Figure 3 (above)**) is misleading in how extensively covered Europe is with chestnut, and purely a consequence of the availability of map resources for European chestnut distribution. The map immediately below it shows very obviously how congruent *D. kuriphilus* and chestnut distributions are based on probability distribution estimates.

### *D. kuriphilus* parasitoid communities

The most prevalent parasitoids attacking *D. kuriphilus* across Europe are cosmopolitan species with broad host ranges (*B. dorsalis, E. urozonus*, *E. brunniventris, Ormyrus pomaceus, Sycophila biguttata, Sycophila variegata, T. flavipes, Torymus geranii*) as well as being some of the most commonly recorded species in oak cynipid galls of the Western Palearctic (Askew *et. al..*, 2013). Some species (*O. pomaceus, T. geranii, S. biguttata, S. variegata, E. urozonus, E. brunniventris*) are Pan-Palearctic, 4 of which have been recorded attacking *D. kuriphilus* in its Eastern Palearctic range (Aebi *et. al.*., 2006). Their cosmopolitan distribution may be indicative of their ability to exploit such a broad repertoire of hosts, and perhaps that at least some of them have a long shared evolutionary history with gallwasp hosts on both *Quercus* and *Castanea*. It is also possible that the identity of these species is erroneous-historically, species that appeared similar were automatically assumed to be the same, regardless of geography, especially when inadequate reference collections were available for comparison. Their status as hyper-generalists may also be inaccurate: many of the abovementioned species are a complex of genetically distinct lineages with highly conserved morphology, or a set of closely related species that are difficult to discern from each other (although identification is still possible). Each morphospecies is therefore not strictly a group of hyper-generalists, but a multitude of species or lineages of varying or unknown host breadths.

### The extent of complexes and cryptic lineages

*E. brunniventris* is a complex of closely related species of oak gall, rose gall and herb gall specialists, but all are commonly recorded from galls other than their main hosts (Askew *et. al..*, 2013). *E. urozonus* is also a complex of many closely related species with similar patterns of ‘leaky’ specialism (Al Khatib *et. al..*, 2014). *B. dorsalis* is composed of two cryptic lineages that cannot be identified morphologically, but show clear ecological differentiation based on the generation of gallwasp hosts that they attack, and by the host oak section of their gallwasp hosts (Nicholls *et. al..*, 2017). In Nearctic communities, the degree of cryptic speciation is particularly enormous in *Sycophila* and *Ormyrus* (*Ormyrus labotus* is putatively composed of 16-18 cryptic lineages) parasitoids of oak galls (Sheikh *et. al.*., 2022; Zhang *et. al..*, 2022), which include a range of broad generalists and specialists of various kinds. What little we know of the gall community in the Eastern Palearctic indicates that many of the gall parasitoids have a broad host range, including gall hosts on oaks across different subgenera, as well as on *D. kuriphilus* on sweet chestnut; however, identification so far has been only by morphotyping due to many undescribed species, and from molecular identification it is clear than none of the current morphotype assignments are representatives of single species, echoing the findings of the Western Palearctic and Nearctic fauna (Stone *et. al.*., unpublished). It has been known for over a decade that the parasitoid *T. flavipes* is composed of at least two cryptic lineages (Kaartinen *et. al..*, 2010), which has been further corroborated by nuclear information by Gil-Tapetado *et. al..* (2022), but as yet nothing is known about their ecology. Given its ubiquity as the most common attacker of both *D. kuriphilus* and *P. quercusilicis* (**Chapter 3**), this topic is explored in greater detail as part of **Chapter 4**.

### Compositional change in parasitoid communities

Across the entire European range, the parasitoid community structure of *D. kuriphilus* varies considerably. Community dissimilarity is high even at local spatial scales of tens of kilometres, and this rapidly increases to a plateau before 1000km predominantly due to increasing levels of turnover and balanced variation (**Figure 4**). This may not be a general trend for all gallwasps as hosts, although there is little data to this end. Bannerman *et. al.*. (2012) found no support of decreasing similarity with increasing geographic distance in the rose gallwasp *Diplolepis variabilis,* noting that generally, communities were incredibly similar in their species compositions. A key difference between *D. kuriphilus* and *D. variabilis* may relate to the relative importance of each host to their parasitoids: parasitoids of *D. kuriphilus* are sourced primarily from a rich community of oak gall hosts scattered amongst a multitude of different oak species, hence a) parasitoids have multiple possible hosts clustered around multiple host trees, b) parasitoids are not obliged to utilise *D. kuriphilus* solely as a host, and c) that this may significantly compartmentalise or fragment local pools of parasitoids centred around any one host species. In contrast, rose galls are generally much less species rich than oak galls and with much less compartmentalisation due to host plant phylogeny (Askew *et. al.*., 2006). Hence high similarity in parasitoid communities of *D. variabilis* might be due to higher degrees of host specificity, or simply to limited availability of alternate hosts.

With the inclusion of sample size and time since establishment as explanatory factors of beta-diversity, only nestedness and gradient components of dissimilarity are significantly affected, and this effect is very minor. The dominant forces of community compositional change are high turnover and balanced variation regardless of sample size and time, which suggests that a significant portion of the parasitoid community attacking *D. kuriphilus* is not fixed on a common set of species, and that over the decades of its European establishment, convergence upon a stable community structure is minimal. All of *D. kuriphilus*’ most common parasitoids are also in the top 20 species most commonly encountered in Europe, both in terms of the number of records, and when considering abundance (Askew *et. al..*, 2013). Hence the assembly of communities appears to be quasi-neutral-in the sense that:

1) The predominant parasitoids of *D. kuriphilus* are largely oak gall parasitoids, so a non-random ecological assemblage (as in a niche based partitioning/ ecological fitting process).
2) Of that assemblage, local communities are largely a function of the probability of detection by the locally most common oak gall generalist parasitoids, hence stochastic with respect to a local pool of ‘ecologically equivalent’ parasitoid species (as in a neutral process).

This conclusion breaks down if most of the common parasitoids are species-complexes (which they are) with distinct ecological partitioning, and otherwise specialised lineages are strongly favouring *D. kuriphilus* as a host! Within the remit of this thesis I was unable to explore this across all species-complexes, but I do elucidate the possibility for lineage sorting amongst *T. flavipes* in **Chapter 4**.

### Parasitoid attack as an effect of increasing establishment period

There are no strong indicators that species richness, or the abundance of parasitoids attacking *D. kuriphilus* are mediated by establishment length. However, most studies provided only single years of data collection. Ideally each locality would have a ‘baseline’ community to compare subsequent years to, and perhaps it would be easier to truly evaluate community development over time this way, instead of viewing data points a given number of years after establishment but at different localities as equivalent. While *D. kuriphilus* has been in Europe for nearly three decades, local populations are nearly all younger than this. Parasitoids are predicted to increase in rates of diversity and attack, owing to 1) the increased probability of detecting a novel host as a function of time, 2) increasing rates of host utility by generalists as a result of adaptations, and 3) the eventual incorporation of novel hosts by specialists adapted to utilise a novel resource (Cornell and Hawkins, 1993). Points 1, 2 and 3 may occur over increasingly longer periods of time: a larger number of *D.* host individuals as a function of range expansions increase the probability of stochastic host ‘collisions’ from a ‘cloud of parasitoid particles’ likely increases the cumulative number of new species recorded to an asymptote, but we would not expect the total number of species or attack rates to increase without some element of selective processes to be invoked.

Cornell and Hawkins (1993) looked at over 80 invader species, with a similar conclusion to this chapter: that generally the diversity and abundance of natural enemies does not increase with establishment time in any meaningful way after accounting for sample size. Perhaps given the current mass of modern literature, it would be possible to answer this question with the restriction that repeated data points be mandatory, although in the case of *D. kuriphilus* in Europe, there is far too little data to be so conservative.

### Final thoughts

The parasitoid community of *D. kuriphilus* within Europe is impressively diverse given its Eastern Palearctic origin and non-oak host. That most of its enemies are oak gallwasp generalist parasitoids (especially when considering abundance) is generally expected, although a significant number of species were non-oak gall members. A paper that missed my initial attention was one of the *D. kuriphilus* community in Madeira, of which 16 of the 19 parasitoid species recorded were non-gall specialists (Aguiar *et. al..*, 2022). Of worthwhile mention is the general lack and or limited diversity of an oak gall community in Madeira. While still dominated in relative abundance by oak gall parasitoids (primarily due to the introduced *T. sinensis*), perhaps the high number of non-gall specialists is an indicator of reduced competition for novel gall resources. Historically community development was considered to be governed predominantly by adaptive processes in the strict evolutionary sense, although ecological fitting is widespread may be the rule rather than the exception, even over evolutionary timescales (Bunnefeld *et. al.*., 2018). However, it is clear that gallwasp parasitoid species have a competitive edge over non-gall specialists, so in their absence, or low presence, they may allow for greater diversity of non-gall specialists such as borer and leaf-miner parasitoids whose host ecologies have significant overlap. At a subfamilial level, parasitoids of leaf miner, borer and non-gallwasp gall specialists are shared, as are many genera (e.g. *Pteromalus, Mesopolobus, Cecidostiba, Torymus, Pediobius, Aprostocetus, Tetrastichus, Baryscapus*).

The influence of *T. sinensis* on parasitoids attacking *D. kuriphilus* is dramatic (Gil-Tapetado *et. al.*., 2021a), and initially I had intended for its distribution to be characterised and included as an explanatory predictor for community developments. However the information on its distribution was far too limited to be included. We know that it does have an influence, so its absence from statistical models are undoubtedly hampered by its exclusion. Variation in the resolution of times of establishment of *D. kuriphilus* also must have added extra noise to the data. Undoubtedly many of the publications’ authors will have more detailed information that would have benefitted this study, but this would have incurred considerably time penalties to the project as to be untenable within the context of the thesis. My second concern was in trying to get a preliminary manuscript submitted to preserve the amount of time and effort that I had invested in synthesising all of these data and then to contact other authors for more detailed information and collaboration. This remains my intention, and I hope that I can expand in much greater depth on the statistical side of this review to make the conclusions more robust. The inclusion of regions as a random effect was a very crude way of accounting for variation in the data, and in a future analysis I will explore the effects of climatic variables on parasitoid communities, which must undoubtedly influence them (Bonsignore *et. al.*., 2020; 2019; Bonsignore and Bernardo, 2018).

