## Supplements 1-3 for "Developments in the European parasitoid community of *Dryocosmus kuriphilus*"

**Appendix 1- Glossary of terms used when compiling European *D. kuriphilus* data.** Some terms also include definitions of their possible options (indicated in green).

| Terms | Definition |
|---|---|
| **Authors** | Authors of the publication |
| **Title** | Title of publication |
| **Year.pub** | Year of publication |
| **Country** | Country where the data was collected. Importantly, if a publication has a multitude of different countries, separate rows are included to account for this |
| **Locality** | Includes more local information about the country where possible. In some cases the resolution is regional, while in others the location is more precise |
| **Lat** | Latitude in decimal format |
| **Lon** | Longitude in decimal format |
| **Lat.lon.origin** | Some papers directly report lat-lon, while others provide less information which requires some manual searching. This column indicates whether the coordinates are generated by me or directly from the paper<br>If a paper reports parasitoid community from combined localities, but provides specific location information, I have produced dummy rows with just the lat lon info included along with a note letting it be known in **Notes.or.errata** |
| 1) Rough centroid | I have located a region on Google Maps and used a rough estimate of that region's centroid to report the coordinates of a location |
| 2) Reported in paper | The coordinate data were directly provided from the paper report |
| **First.record** | The year in which *D. kuriphilus* was first recorded at a location. Where possible, this measure is as accurate as can be (regional-level), though in a few cases this indicates establishment at a country-level resolution because it is the only information available. |
| **Year.study** | The year(s) in which a study took place. Some papers have multiple years of studies, but only provide data collectively. In this case, a range of years is reported in a single row e.g. 2013-2015 (see also **Length.of.study.period**) |
| **Years.since.first.record** | **Year.study** minus the **First.record.** In cases where multiple years of data are recorded within a single data entry as indicated in year of study, the last year of the study is used. |
| **Length.of.study.period** | The increment of time that a row refers to. Usually this will be a single year, but in cases with a range it is the number of years in that range (see **Year.study**) |
| **Report.format** | The format that parasitoid records are reported in this excel file. As much as possible, the data is reported as it is from the original source, but on some occasions, the data is too incomplete to provide better information beyond presence/ absence |
| 1) New species- number.of.individuals | Only applies to one paper from Greece with the discovery of *Synergus kuriphili* . It did not include any other rearing info |
| 2) Percentage of emerged parasitoids | The percentage of parasitoids reared from galls. This is NOT the percentage of parasitoids as a proportion of all insects emerged (i.e. including DK) |
| 3) Number of parasitoids | Raw count of parasitoids |
| 4) Presence | Expressed as an X |
| **Notes.or.errata** | Outlines any random information that may be worth noting, or errors in the paper. As much as possible I have tried to create columns outlining important information in a less anecdotal way |
| **Fresh.vs.winter.rearings** | Indicates whether rearings include fresh and or overwintered galls within rearings |
| 1) Not stated | Not reported or ambiguous |
| 2) Fresh | Fresh galls collected |
| 3) Winter | Overwintered galls collected |
| 4) Fresh and overwintering | Fresh and overwintered galls collected |
| **No.of.galls** | Self explanatory, BUT, there may be some instances when, for example, detailed year-by-year information is given for parasitoids, but the number of galls is a collated number across years. In that case, I have produced a single dummy row indicating the number of galls for the entirety of the study period, and no additional information on parasitoids in that row |
| **No.of.spp** | This includes taxa that are not resolved to species-level i.e generic level. Hence, this could mean the number of taxa when uncertainty exists. |

**Appendix 2. The number of localities sampled for *D. kuriphilus* parasitoids in its European range.** The number of localities include information about with multiple years of data collection, given in raw numbers in addition to the percentage total for each country. The total number of localities is reported the far right of the graph along with the percentage of localities as a proportion of the European total.

| Country | Title | Yr. pub | Authors | N. localities | N. yrs. data | N. galls | | |
|---|---|---|---|---|---|---|---|---|
| Italy | Parasitoid Recruitment to the Globally Invasive Chestnut Gall Wasp Dryocosmus kuriphilus | 2006 | Aebi et al | 1 | 4 | 11113 | * | |
| Italy | Endemic Parasitoids of Dryocosmus kuriphilus Yasumatsu (Hymenoptera: Cynipidae) in Central Italy | 2009 | Speranza et al | 1 | 1 | - | | |
| Italy | New association between Dryocosmus kuriphilus and Torymus flavipes in chestnut trees in the Bologna area (Italy): first results | 2011 | Santi and Maini | 2 | 2 | 1000 | | |
| Italy | Orthopelma mediator (THUNBERG) (Hymenoptera: Ichneumonidae) and the native parasitoid complex of Dryocosmus kuriphilus YASUMATSU (Hymenoptera: Cynipidae) in Lombardy (Italy) | 2013 | Boriani et al | 1 | 6 | - | | |
| Croatia | Recruitment of native parasitoids to a new invasive host: first results of Dryocosmus kuriphilus parasitoid assemblage in Croatia | 2013 | Matosevic and Melika | 4 | 2 | 20598 | | |
| Italy | Native parasitoids associated with Dryocosmus kuriphilus in Tuscany, Italy | 2013 | Panzavolta et al | 3 | 2 | 2276 | | |
| Italy | Chalcid parasitoid community associated with the invading pest Dryocosmus kuriphilus in north-western Italy | 2013 | Quacchia et al | 1 | 5 | 415224 | † | ** |
| Turkey | First reports on the natural enemy fauna of the chestnut gallwasp, Dryocosmus kuriphilus Yasumatsu (Hymenoptera: Cynipidae) in Yalova, Turkey | 2014 | Doganlar | 5 | 1 | 1154 | | |
| Italy | Indigenous parasitoids associated with Dryocosmus kuriphilus in a chestnut production area of Emilia Romagna (Italy) | 2015 | Francati et al | 1 | 3 | 12015 | | |
| Slovenia | Invasive chestnut gall wasp Dryocosmus kuriphilus (Hymenoptera: Cynipidae), its native parasitoid community and association with oak gall wasps in Slovenia | 2015 | Kos et al | 1 | 4 | 36948 | ** | *** |
| Slovenia | Invasive chestnut gall wasp Dryocosmus kuriphilus (Hymenoptera: Cynipidae), its native parasitoid community and association with oak gall wasps in Slovenia | 2015 | Kos et al | 1 | 3 | - | ** | *** |
| Slovenia | Invasive chestnut gall wasp Dryocosmus kuriphilus (Hymenoptera: Cynipidae), its native parasitoid community and association with oak gall wasps in Slovenia | 2015 | Kos et al | 2 | 1 | - | ** | *** |
| Italy | Survey of indigenous parasitoids affecting the invasive chestnut gall wasp Dryocosmus kuriphilus in the Friuli Venezia Giulia region (North-East Italy) | 2016 | Bernadinelli et al | 1 | 1 | - | | |
| Italy | Native and introduced parasitoids in the biocontrol of Dryocosmus kuriphilus in Veneto (Italy) | 2016 | Colombari and Batisti | 1 | 3 | 6415 | | |
| Italy | Native and introduced parasitoids in the biocontrol of Dryocosmus kuriphilus in Veneto (Italy) | 2016 | Colombari and Batisti | 2 | 2 | 4790 | | |
| Italy | Native and introduced parasitoids in the biocontrol of Dryocosmus kuriphilus in Veneto (Italy) | 2016 | Colombari and Batisti | 20 | 1 | 9442 | | |
| Italy | Studies on the remnants of the parasitoid larvae of Dryocosmus kuriphilus Yasumatsu (Hymenoptera: Cynipidae), in the galls collected in two places in Italy for determining parasitism level and type of parasitoids | 2017 | Doganlar | 2 | 1 | 300 | | |
| Spain | Dryocosmus kuriphilus Yasumatsu, 1951 (Hymenoptera: Cynipidae) in Galicia (NW Spain): pest dispersion, associated parasitoids and first biological control attempts | 2017 | Pérez-Otero et al | 1 | 4 | 18860 | | |
| Italy | Do Torymus sinensis (Hymenoptera: Torymidae) and agroforestry system affect native parasitoids associated with the Asian chestnut gall wasp? | 2018 | Ferracini et al | 34 | 1 | 17000 | | |
| Portugal | Native parasitoids associated with Dryocosmus kuriphilus Yasumatsu in Portugal: main species and parasitism rates | 2018 | Lobo Santos et al | 1 | 4 | 5760 | | |
| Portugal | Native parasitoids associated with Dryocosmus kuriphilus Yasumatsu in Portugal: main species and parasitism rates | 2018 | Lobo Santos et al | 1 | 1 | 1440 | | |
| Greece | New species of cynipid inquiline, Saphonecrus kuriphilusi (Hymenoptera: Cynipidae: Synergini), from Drycocosmus kuriphilus galls in Greece | 2018 | Melika et al | 1 | 1 | 1 | *** | † |
| Italy | Population Dynamics of Native Parasitoids Associated with the Asian Chestnut Gall Wasp (Dryocosmus kuriphilus) in Italy | 2018 | Panzavolta et al | 2 | 3 | 1680 | | |
| Italy | Tracking seasonal emergence dynamics of an invasive gall wasp and its associated parasitoids with an open-source, microcontroller-based device | 2018 | Rondoni et al | 1 | 2 | 800 | | |
| Spain | Los enemigos naturales de la avispilla asiática del castaií.o (Dryocosmus kuriphilus) en Galicia | 2018 | Santolamazza-Carbone | 8 | 1 | 2632 | | |
| Italy | Effects of environmental parameters on the chestnut gall wasp and its complex of indigenous parasitoids | 2018 | Bonsignore et al | 1 | 3 | - | ‡ | ‡ |
| Italy | Environmental thermal levels affect the phenological relationships between the chestnut gall wasp and its parasitoids | 2019 | Bonsignore et al | 3 | 2 | - | | |
| Italy | Environmental thermal levels affect the phenological relationships between the chestnut gall wasp and its parasitoids | 2019 | Bonsignore et al | 1 | 1 | - | | |
| Spain | The invasive ACGW Dryocosmus kuriphilus (Hymenoptera: Cynipidae) in Spain: native parasitoid recruitment and association with oak gall inducers in Catalonia | 2019 | Jara-Chiquito et al | 7 | 4 | 19445 | | |
| France | The open bar is closed: restructuration of a native parasitoid community following successful control of an invasive pest | 2019 | Muru et al | 1 | 5 | 284425 | | |
| Italy | Short-Term Cold Stress Affects Parasitism on the Asian Chestnut Gall Wasp Dryocosmus kuriphilus | 2020 | Bonsignore et al | 1 | 2 | 7423 | | |
| Spain | Characterization of native parasitoid community associated with the invasive pest Dryocosmus kuriphilus (Hymenoptera: Cynipidae) in Cantabria (Northern Spain) | 2020 | Dorado et al | 2 | 1 | 500 | | |
| Spain | Differences in native and introduced chalcid parasitoid communities recruited by the invasive chestnut pest Dryocosmus kuriphilus in two Iberian territories | 2020 | Gil-Tapetado et al | 15 | 3 | 18074 | | |
| Spain | Differences in native and introduced chalcid parasitoid communities recruited by the invasive chestnut pest Dryocosmus kuriphilus in two Iberian territories | 2020 | Gil-Tapetado et al | 1 | 2 | - | | |
| Spain | Newly invaded territories by Dryocosmus kuriphilus in Spain and first records of Torymus sinensis in the Sistema Central | 2020 | Gil-Tapetado et al | 2 | 1 | 305 | | |
| England | Recruitment of native parasitoids by an introduced gall wasp Dryocosmus kuriphilus Yasumatsu, 1951 (Hymenoptera: Cynipidae) in Britain and France | 2020 | Jennings and Askew | 2 | 1 | - | | |
| France | Recruitment of native parasitoids by an introduced gall wasp Dryocosmus kuriphilus Yasumatsu, 1951 (Hymenoptera: Cynipidae) in Britain and France | 2020 | Jennings and Askew | 1 | 4 | - | | |
| Croatia | Diversity and surge in abundance of native parasitoid communities prior to the onset of Torymus sinensis on the Asian chestnut gall wasp (Dryocosmus kuriphilus) in Slovenia, Croatia and Hungary | 2020 | Kos et al | 1 | 3 | 22589 | | |
| Slovenia | Diversity and surge in abundance of native parasitoid communities prior to the onset of Torymus sinensis on the Asian chestnut gall wasp (Dryocosmus kuriphilus) in Slovenia, Croatia and Hungary | 2020 | Kos et al | 1 | 4 | 50322 | | |
| Hungary | Diversity and surge in abundance of native parasitoid communities prior to the onset of Torymus sinensis on the Asian chestnut gall wasp (Dryocosmus kuriphilus) in Slovenia, Croatia and Hungary | 2020 | Kos et al | 1 | 3 | 16655 | | |
| Spain | Gall size of Dryocosmus kuriphilus limits down-regulation by native parasitoids | 2021 | Gil-Tapetado et al | 5 | 1 | 12525 | | |
| Italy | Disentangling the effects of the invasive pest, Dryocosmus kuriphilus, and the introduction of the biocontrol agent, Torymus sinensis, on native parasitoids in an isolated insular chestnut-growing area | 2021 | Loru et al | 16 | 9 | 12525 | | |
| Italy | Julja Ernst thesis data | Unpublished | Ernst | 6 | 1 | 4943 | | |
| Italy | Julja Ernst thesis data | Unpublished | Ernst | 6 | 1 | 3006 | | |
| Italy | Julja Ernst thesis data | Unpublished | Ernst | 3 | 1 | 3000 | | |
| Switzerland | Julja Ernst thesis data | Unpublished | Ernst | 1 | 2 | - | | |
| Switzerland | Julja Ernst thesis data | Unpublished | Ernst | 1 | 5 | - | | |
| England | Fera data | Unpublished | Malumphy | 1 | 4 | - | | |
| England | Fera data | Unpublished | Malumphy | 7 | 1 | - | | |
| England | My data | Unpublished | McCormack | 11 | 1 | 4400 | | |

* No. of galls not recorded in the first year of the study
** No. of Sites reduced to one community but there were actually many
*** No. of galls for all sites reported as single figure
† Description of new species only
‡ No. of Sites reduced to one community but there were actually 3

**Appendix 3. The relative proportion of the top 20 parasitoid species of *D. kuriphilus* across Europe (below) and the number of parasitoids reared (top) with the inclusion of all *Eupelmus* morphospecies.** Data from Ferracini *et. al.*. (2017) are excluded (see **Methods**)

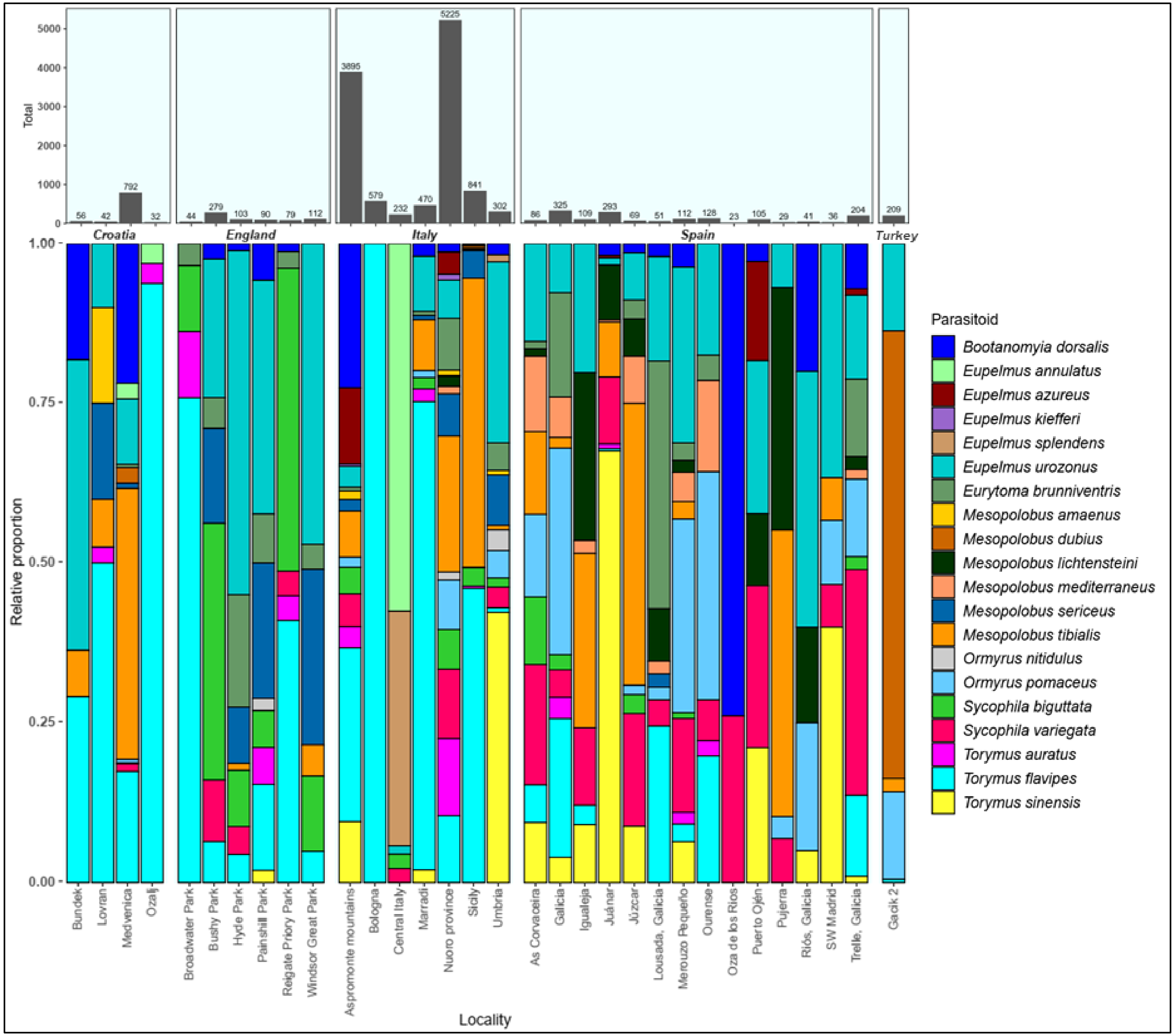
